# A Non-Canonical IRAK Signaling Pathway Triggered by DNA Damage

**DOI:** 10.1101/2023.02.08.527716

**Authors:** Yuanyuan Li, Richa B. Shah, Samanta Sarti, Alicia L. Belcher, Brian J. Lee, Andrej Gorbatenko, Francesca Nemati, Ian Yu, Zoe Stanley, Zhengping Shao, Jose M. Silva, Shan Zha, Samuel Sidi

**Affiliations:** Department of Medicine, Division of Hematology and Medical Oncology, Tisch Cancer Institute, Icahn School of Medicine at Mount Sinai, New York, NY; Department of Cell, Developmental and Regenerative Biology, The Graduate School of Biomedical Sciences, Icahn School of Medicine at Mount Sinai, New York, NY; Institute for Cancer Genetics, College of Physicians and Surgeons, Columbia University, New York, NY; Department of Oncological Sciences, Icahn School of Medicine at Mount Sinai, New York, NY; Department of Pathology, Icahn School of Medicine at Mount Sinai, New York, NY; Department of Medical Biochemistry, Amsterdam UMC, University of Amsterdam, The Netherlands; Division of Pediatric Oncology, Hematology and Stem Cell Transplantation, Department of Pediatrics, College of Physicians and Surgeons, Columbia University, New York, NY

## Abstract

Interleukin-1 receptor (IL-1R)-associated kinases (IRAKs) are core effectors of Toll-like receptor (TLR) and IL-1R signaling, with no reported roles outside of innate immunity. We find that vertebrate cells exposed to ionizing radiation (IR) sequentially activate IRAK4 and IRAK1 through a phosphorylation cascade mirroring that induced by TLR/IL-1R, resulting in a potent anti-apoptotic response. However, IR-induced IRAK1 activation does not require the receptors or the IRAK4/1 adaptor protein MyD88, and instead of remaining in the cytoplasm, the activated kinase is immediately transported to the nucleus via a conserved nuclear localization signal. We identify: double-strand DNA breaks (DSBs) as the biologic trigger for this pathway; the E3 ubiquitin ligase Pellino1 as the scaffold enabling IRAK4/1 activation in place of TLR/IL-1R-MyD88; and the pro-apoptotic PIDDosome (PIDD1-RAIDD-caspase-2) as a critical downstream target in the nucleus. The data delineate a non-canonical IRAK signaling pathway derived from, or ancestral to, TLR signaling. This DSB detection pathway, which is also activated by genotoxic chemotherapies, provides multiple actionable targets for overcoming tumor resistance to mainstay cancer treatments.

## Introduction

Interleukin-1 receptor (IL-1R)-associated kinase 1 (IRAK1) is an evolutionarily conserved death domain (DD)-containing protein kinase whose *Drosophila* homolog, pelle, transduces dorso-ventral patterning and microbial cues recognized by the transmembrane receptor, Toll (Gay and Keith, 1991; Hashimoto et al., 1988; Janssens and Beyaert, 2003; Lemaitre et al., 1996; Nusslein-Volhard, 2022; Shelton and Wasserman, 1993). The immune function was found to be conserved in vertebrates, in which the kinase transduces pathogen cues recognized by Toll-like receptors (TLR) and IL-1R (Cao et al., 1996; Flannery and Bowie, 2010; Medzhitov et al., 1997; O’Neill et al., 2013). As in flies, TLR/IL-1R-induced IRAK1 activation culminates in the activation of pro-inflammatory signaling cascades including NF-κB, p38/MAPK and JNK (Kawasaki and Kawai, 2014; O’Neill et al., 2013), overall defining a core branch of innate immunity across species (Fitzgerald and Kagan, 2020; Lemaitre, 2004).

The mechanism of IRAK1 activation by TLR/IL-1R was elucidated by several groups who identified Myeloid Differentiation Primary Response 88 (MyD88) (Lord et al., 1990), a protein of previously unknown function (Deguine and Barton, 2014), as the key adaptor molecule responsible for the recruitment of IRAK1 and its sister kinase, IRAK4, to the ligated receptors (Burns et al., 1998; Medzhitov et al., 1998; Muzio et al., 1997; Wesche et al., 1997). MyD88 harbors a Toll/IL-1R homology (TIR) domain and a DD which respectively bind the receptor and IRAK kinases through homotypic TIR:TIR and DD:DD interactions, thus physically bridging the molecules. This ultimately results in the formation of the MyDDosome complex (MyD88-IRAK4-IRAK1) at the inner surface of the cell (Latty et al., 2018; Lin et al., 2010; Motshwene et al., 2009; Wang et al., 2017). The MyDDosome provides the necessary activation platform for IRAKs: Only once in the MyDDosome can IRAK4 dimerize and *trans*- autophosphorylate, resulting in its activation (Cheng et al., 2007; Cushing et al., 2014; Ferrao et al., 2014; Wang et al., 2017). This proximity-induced activation of IRAK4 is the key initiating step in IRAK1 activation, with most (Cushing et al., 2014; Ferrao et al., 2014; Kollewe et al., 2004; Latty et al., 2018; Wang et al., 2017; Wang et al., 2006) but not all (Vollmer et al., 2017) models implicating IRAK4-mediated phosphorylation of IRAK1 on residue T209 as responsible for IRAK1 activation. IRAK1 then fully activates via autophosphorylation on T387 in the activation loop and, in turn, dissociates from the MyDDosome (Kollewe et al., 2004; Wang et al., 2017). To date, no pathway other than TLR/IL-1R has been reported to signal through IRAK1/4, and while TLRs can signal independently of MyD88, all TLR/IL-1R pathways that signal through IRAKs do so through MyD88 (Fitzgerald and Kagan, 2020; O’Neill et al., 2013).

The notion that IRAK kinases are confined to TLR/IL-1R signaling was recently challenged when an unbiased screen in zebrafish identified IRAK1 as essential for cell survival in response to ionizing radiation (IR) (Liu et al., 2019). This pro-survival signaling function is conserved in human cells, in which it appears to drive intrinsic tumor resistance to radiation therapy (R-RT). Rather than signaling through NF-κB, p38/MAPK, JNK or ERK, IRAK1 was found to act, at least in part, by preventing IR-induced apoptosis mediated by the PIDDosome complex (PIDD-RAIDD-caspase-2) (Tinel and Tschopp, 2004). Most surprisingly, while IR-induced IRAK1 signaling appeared to involve IRAK4, it did not appear to require MyD88 (Liu et al., 2019). These observations suggested that IRAK1 might serve in a pathway distinct from the canonical, TLR/IL-1R axis. Here, we present evidence in support of such a non-canonical pathway of IRAK1 signaling, which is responsible for sensing and transducing DNA damage into an anti-apoptotic response. Key distinguishing features include a distinct IRAK1 activation platform operating in the cytoplasm, not the cell surface, and distinct downstream signaling which requires rapid transport of the activated kinase from the cytoplasm into the nucleus.

## Results

### IR-induced IRAK1 signaling requires IRAK4 but not MyDDosome assembly

RNA interference (RNAi) studies in HeLa cells suggested that IRAK4, like IRAK1, is required for cell survival in response to IR and might function in the same, MyD88-independent pathway (Liu et al., 2019). We confirmed these RNAi results in additional cell lines (Figure S1A-B) and in an *IRAK4* knockout line obtained by CRISPR/Cas9 editing (Figure 1A-B; and Figure 1C, bars 2 vs. 4). The radiosensitive phenotype of *IRAK4^—/—^* cells was rescued by wild-type (WT) but not kinase dead (D329) FLAG-IRAK4 (Figures 1C and S1C) (De et al., 2018). These results were recapitulated in zebrafish *p53^M214K/M214K^* (*p53^MK/MK^*) mutant embryos, a whole-animal model of tumor R-RT (Sidi et al., 2008) in which IR-induced IRAK1 signaling was originally described (Liu et al., 2019). Morpholino antisense oligonucleotide (MO)-mediated knockdown of Irak4, similar to that of Irak1 (Liu et al., 2019), strongly sensitized the mutant fish to IR, as evidenced by a marked uptake of the cell death marker acridine orange in developing spinal cords (Figure 1D, compare panels (2) and (4), and Figures 1E-G). These effects were on-target because a standard control MO (std MO) had no effect and because Irak4-depleted fish were rescued by co-injection with human IRAK4 mRNA, but not IRAK4^D329A^ mRNA (Figure 1D, compare panels (4), (6) and (10), and Figures 1E,G). These experiments demonstrated an evolutionarily conserved pro-survival role for IRAK4 in irradiated cells and that this function requires IRAK4 catalytic activity, similar to previously observed for IRAK1 (Liu et al., 2019).

**Figure 1.**
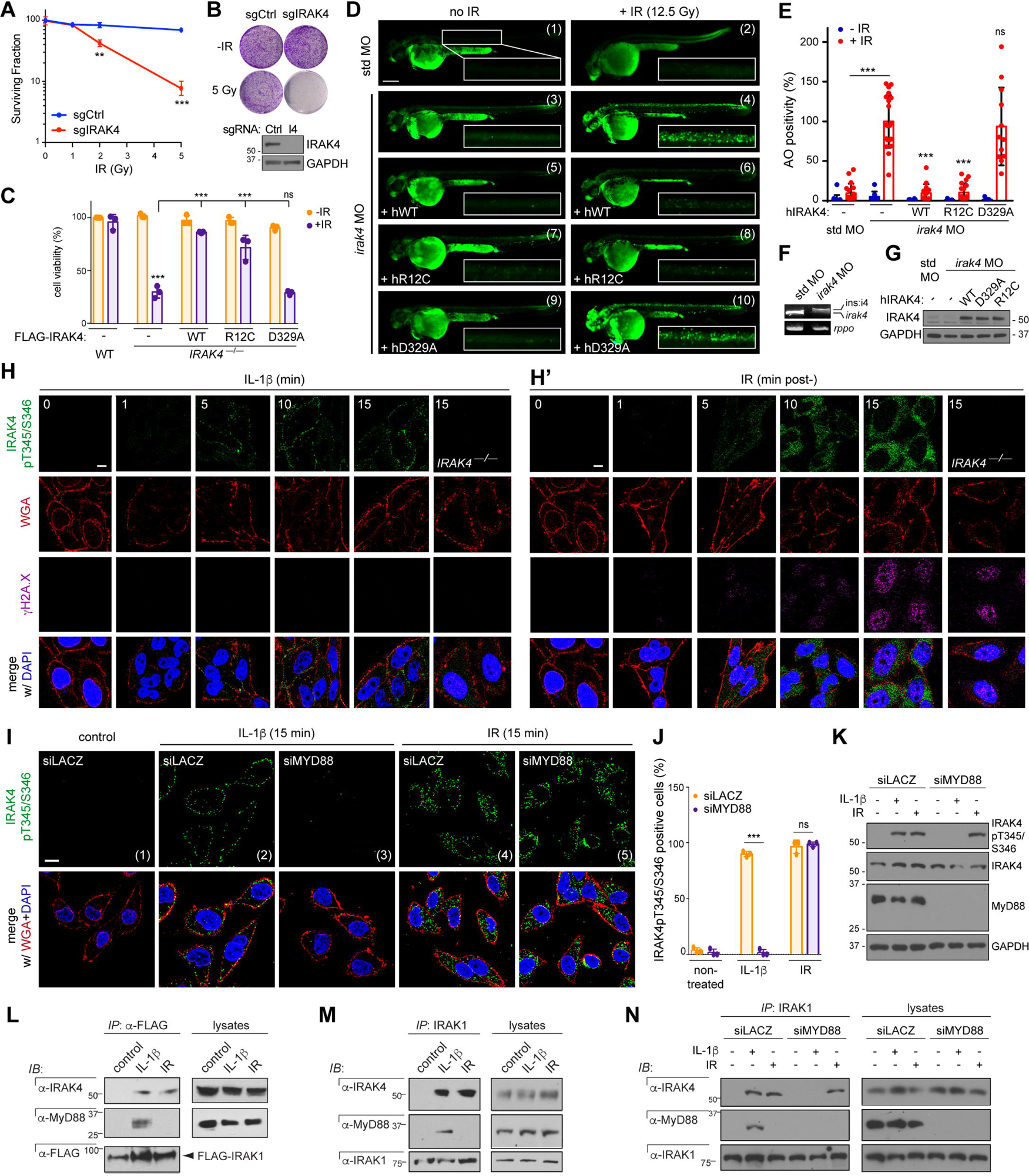
IRAK4 associates with IRAK1 and is required for cell survival after IR in a MyD88-independent manner. (A-B) Parent and CRISPR/Cas9-generated *IRAK4^—/—^* HeLa cells were treated with indicated doses of X-IR and colonies were counted (A) and imaged (B) at 12 days post-IR. Data are means +/− SD of 3 independent experiments with \*\**p* < 0.01 and \*\*\**p* < 0.001, two-tailed Student’s t-test. Western blot verifying IRAK4 knockout also shown in (B). (C) Parent and *IRAK4^—/—^* cells reconstituted with indicated FLAG-IRAK4 constructs were treated with or without IR (7.5 Gy). Cells were stained with the vital dye alamarBlue 72 hr post- IR. Data are means +/− SD of 3 independent experiments. \*\*\**p* < 0.0001, two-tailed Student’s t-test. Uniform expression levels of FLAG-IRAK4 variants shown in Figure S1C. (D) *p53^MK/MK^* zebrafish embryos were co-injected at the 1 cell stage with the indicated MOs with or without synthetic mRNAs for the indicated human (h) FLAG-IRAK4 variants (hWT, hR12C and h329A). Embryos were treated with or without whole-body IR (12.5 Gy) 18 hours later and stained with acridine orange (AO) at 24 hpIR to label apoptotic cells in vivo. Spinal cord areas dorsal to the yolk tube (boxed in each panel) were used for quantification shown in (E). Scale bar, 0.2 mm. (E) Quantification of spinal cord areas from (D) of at least 6 embryos per condition over 3 independent experiments. Data are means +/− SEM, \**p* < 0.05, \*\**p* < 0.01, \*\*\**p* < 0.001, two-tailed Student’s t-test. (F) mRNA extracts from embryos injected with indicated MOs were analyzed by RT-PCR for *irak4* and *rppo*. Note that the *irak4* MO, which targets the exon4/intron4 splice junction, results in the retention of intron 4 (199 bp) in the mature mRNA, as verified by sequencing. (G) Pooled whole-embryo extracts from (D) analyzed by western blot to verify uniform expression levels of human IRAK4 constructs. (H, H’) HeLa cells grown on cover slips were treated with IL-1β (0.1 μg/mL) (G) or IR (7.5 Gy) (G’), fixed at indicated time points (min) after treatment, stained with indicated antibodies and co-stained with membrane marker wheat germ agglutinin (WGA) and nuclear marker DAPI. Confocal images representative of *n*=3 independent experiments. Scale bar, 10 μm. (I) HeLa cells transfected with indicated siRNA were grown on cover slips, treated with or without IL-1β (0.1 μg/mL) or IR (7.5 Gy), fixed 15 min after treatment, stained with IRAK4pT345/S346 antibody and co-stained with WGA and DAPI. Confocal images representative of *n*=3 independent experiments were quantified in (I). Single channels shown in Figure S1D. Scale bar, 10 μm. (J) Quantification of IRAK4pT345/S346 stains such as in (H). Data are means +/− SD of 3 independent experiments. \*\*\**p* < 0.0001; ns, non-significant; two-tailed Student’s t-test. (K) Whole-cell lysates from cells in (H) were analyzed by western blot with indicated antibodies. (L-M) FLAG-IRAK1-transfected (K) or non-transfected (L) HeLa cells were treated with or without IL-1β (0.1 μg/mL) or IR (7.5 Gy) and lysed 15 min post-treatment. FLAG (K) or endogenous IRAK1 (L) immunoprecipitates were analyzed by western blot with indicated antibodies. *IP*, immunoprecipitate; *IB*, immunoblot. (N) HeLa cells transfected with the indicated siRNAs were treated with or without IL-1β (0.1 μg/mL) or IR (7.5 Gy) and lysed 15 min post-treatment. IRAK1 immunoprecipitates were analyzed by western blot with indicated antibodies.

Should IRAK4 function in the same pathway as IRAK1, then IR would be expected to activate IRAK4 in a MyD88-independent manner (Liu et al., 2019). Most models of IRAK4 activation involve dimerization-induced, IRAK4 *trans-*autophosphorylation on multiple sites encompassing T345 and S346 (Cheng et al., 2007; Cushing et al., 2014; Ferrao et al., 2014). We tested an antibody to IRAK4pT345/S346 in immunofluorescence assays. We first analyzed IL-1β-treated cells, which would be expected to activate IRAK4 at the cell surface via MyDDosome formation at ligated IL-1Rs. The anti-IRAK4pT345/S346 signals matched this prediction: signals were detected at the inner cell periphery from 5 min post-treatment on (Figure 1H) and were absent in *IRAK4^—/—^* cells (Figure 1H) and MyD88-depleted cells (Figure 1I, compare panels (2) and (3), and Figure 1J). These experiments identified the IRAK4pT345/S346 antibody as a reliable marker of IRAK4 activation.

In response to IR, IRAK4 was activated with similar kinetics, with IRAK4pT345/S346 signals first detected as early as 5 min post-IR and peaking at 15 min (Figure 1H’). However, in sharp contrast with IL-1β, IR led to IRAK4 activation not at the cell surface but in the cytoplasm (Figure 1H’), and while the signal was absent in *IRAK4^—/—^* cells, it was unaffected by depletion of MyD88 (Figure 1I, compare panels (4) and (5); Figures 1J-K; and Figure S1D). The MyD88-independence of IR-induced IRAK4 activation was further confirmed by immunoblot (Figure 1K). Together, the cytoplasmic localization of IR-induced IRAK4 autophosphorylation and its non-reliance on MyD88 provided first independent support for the existence of an alternate, MyDDosome-independent route to IRAK1/4 activation in human cells.

To verify this biochemically, we performed a series of co-immunoprecipitation assays. While IRAK1 associated with both MyD88 and IRAK4 in response to IL-1β, as expected, it associated with IRAK4 but not MyD88 after IR (Figure 1L; endogenous pulldowns shown in Figure 1M). Additionally, while MyD88 was required for the IRAK4-IRAK1 interaction in IL-1β- treated cells, again as expected, it was not after IR (Figure 1N). Consistent with these data, an IRAK4 mutant unable to bind MyD88, IRAK4^R12C^ (De et al., 2018), rescued irradiated *IRAK4^—/—^* cells and Irak4-depleted zebrafish as effectively as IRAK4^WT^ (Figure 1C-E). Thus, while kinetically similar, activation of IRAK4 by IR differs from that by IL-1β both spatially (cytoplasm vs. cell surface) and mechanistically (MyDDosome-independent vs. dependent).

### Active IRAK1 accumulates in the nucleus of irradiated cells

Given the unexpected cytoplasmic localization of IR-induced IRAK4 activation, we sought to investigate that of its presumptive substrate, IRAK1, and product, IRAK1pT209, in irradiated cells. To identify reliable markers, we again trialed several antibodies in IL-1β-treated cells, in which both the native and phosphorylated species should be detected at the inner cell surface within minutes of treatment. Antibodies to IRAK1 and IRAK1pT209 were identified which produced the expected IL-1β-induced cell surface signals (Figure 2A-B, and note the absence of signals in *IRAK1^—/—^* cells).

**Figure 2.**
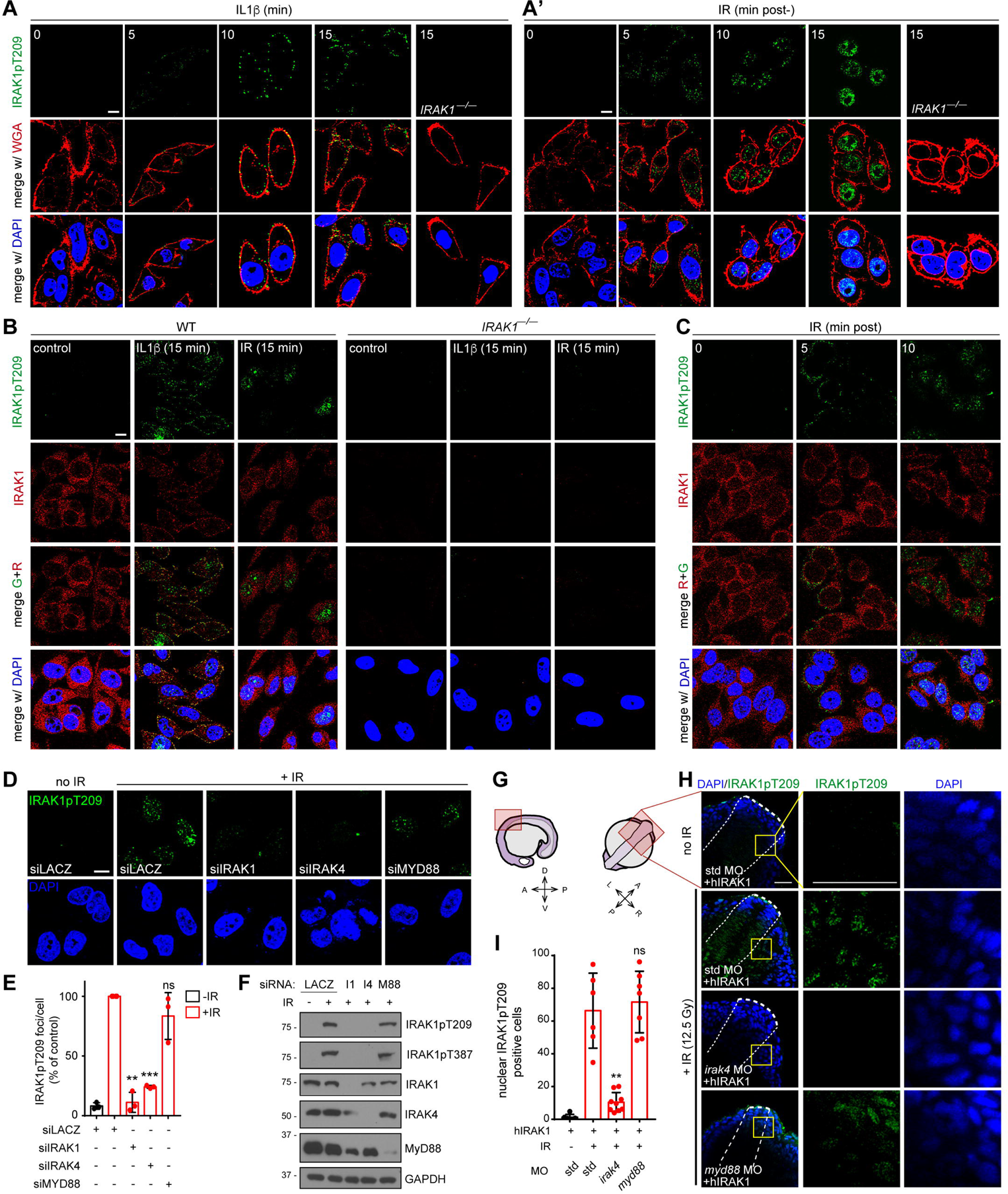
Active IRAK1 accumulates in the nucleus of irradiated cells in an IRAK4- dependent but MyD88-independent manner. (A-A’) Parental and *IRAK1^—/—^* HeLa cells grown on cover slips were treated with IL-1β (0.1 μg/mL (A) or IR (7.5 Gy) (A’), fixed at indicated time points (min) after treatment, stained with IRAK1pT209 antibody and co-stained with membrane marker WGA and DAPI. Confocal images representative of 3 independent experiments. (B) Parental and *IRAK1^—/—^* HeLa cells grown on cover slips were treated with IL-1β (0.1 μg/mL) or IR (7.5 Gy), fixed at 15 min, stained with indicated antibodies and co-stained with DAPI. Confocal images representative of 3 independent experiments. (C) HeLa cells grown on cover slips were treated with IL-1β (0.1 μg/mL) or IR (7.5 Gy), fixed at indicated time points, stained with indicated antibodies and co-stained with DAPI. Confocal images representative of 3 independent experiments. (D) HeLa cells transfected with indicated siRNA were grown on cover slips, treated with or without IR (7.5 Gy), fixed 15 min after treatment, stained with IRAK1pT209 antibody and co-stained with DAPI. Confocal images representative of *n*=3 independent experiments were quantified in (E). (E) Quantification of IRAK1pT209 stains such as in (D). Data are means +/− SD of 3 independent experiments. Statistical significance vs. bar 2: \*\**p*<0.01, \*\*\**p* < 0.001; two-tailed Student’s t-test. (F) Cells as in (D) were lysed and analyzed by western blot with indicated antibodies. I1, IRAK1; I4, IRAK4; M88, MYD88. (G) Schematic of lateral and dorsal views of the 18-hour zebrafish embryo, with imaged area (see H) boxed in pink. A, anterior; P, posterior; V, ventral; D, dorsal; L, left; R, right. (H) Zebrafish *p53^MK/MK^* mutant embryos were co-injected at the 1 cell stage with the indicated MOs with or without synthetic mRNA for human FLAG-IRAK1 (hIRAK1) to detect IRAK1p209 in vivo (the IRAK1pT209 antibody does not cross-react with zebrafish Irak1). Embryos were treated with or without whole-body IR (12.5 Gy) 18 hours later, fixed at 15 min post-IR, and stained as whole-mounts with IRAK1pT209 antibody and DAPI. Higher magnification images of boxed areas shown to the right were used for quantification shown in (I). Anterior and spinal cord areas delineated by thick and thin dashed lines, respectively. Scale bar, 40 μm. (I) Quantification of spinal cord areas (such as boxed in (H)) of at least 6 embryos per condition over 3 independent experiments. Data are means +/− SEM, statistical significance vs. bar 2: \*\**p* < 0.01, two-tailed Student’s t-test.

In response to IR, native IRAK1 was detected in the cytoplasm, not at the cell surface, and remained cytoplasmic throughout (Figures 2B and S2A). IRAK1pT209 was first detected in the cytoplasm at 5 min post-IR (Figures 2A’ and S2B), which was consistent with the localization of active IRAK4 and native IRAK1 in irradiated cells (Figures 1G and S2A). Strikingly however, by 15 min, IRAK1pT209 was detected exclusively in the nucleus (Figures 2A’ and S2B), in sharp contrast with its cell-surface localization in IL-1β-treated cells (Figure 2A). The IR-induced nuclear localization of IRAK1pT209 was confirmed in multiple cell lines (Figure S2C-D) and was also observed in irradiated zebrafish (Figure S2E).

Irradiated cells analyzed between the 5- and 15- min timepoints showed varying levels of cytoplasmic and nuclear IRAK1pT209 foci. At 10 min, the activated kinase could be found both in the cytoplasm and nucleus (Figures S2B) or already exclusively in the nucleus (Figure 2A’). Double-staining with IRAK1 and IRAK1pT209 antibodies confirmed that the native kinase remained in the cytoplasm as the nuclear IRAK1pT209 signal progressively increased (Figures 2B-C). These observations provided first evidence of a possible transport of activated IRAK1 into the nucleus (see Figure 4 below). Nuclear accumulation of IRAK1pT209 peaked at 15 min, was retained for 6 hours, and declined by 12-24h (Figure S2F-G).

Consistent with residue T209 being the target of IRAK4, IR-induced IRAK1pT209 signals were undetectable in IRAK4-depleted cells (Figure 2D), *IRAK4^—/—^* cells (Figure S2H) and Irak4- depleted zebrafish embryos (Figure 2G-I). WT IRAK4 restored T209 phosphorylation in irradiated *IRAK4^—/—^* cells while kinase-dead IRAK4^R329A^ failed to do so, confirming the requirement for IRAK4 catalytic activity (Figure S2H-I). In contrast to IRAK4, MyD88 was not required for IRAK1 activation in either human or zebrafish cells (Figures 2D-H). IRAK4^R12C^, which fails to bind MyD88 (De et al., 2018), restored IR-induced IRAK1 T209 phosphorylation in *IRAK4^—/—^* cells as efficiently as the WT kinase (Figure S2H-I). Collectively, these results revealed the existence of an evolutionarily conserved, MyD88-independent route to IRAK1 activation by IRAK4. This distinct IRAK signaling axis can be activated by IR and operates with similar kinetics to the canonical, TLR/IL-1R pathway, but in a spatially distinct manner.

### IR-induced IRAK1 activation triggers its autophosphorylation in the nucleus

Biochemical and structural evidence suggests that once phosphorylated by IRAK4 on T209, IRAK1 autophosphorylates on T387 within the activation loop to achieve full activation (Kollewe et al., 2004; Wang et al., 2017). A commercially available antibody raised against IRAK1pT387 produced nuclear signals in irradiated cells reminiscent of the IRAK1pT209 stains shown above (Figures 3A-B, 3I and 3J; five additional cell lines shown in Figure S3A). IR- induced nuclear IRAK1pT387 foci were absent in *IRAK1^—/—^* cells and were restored via transfection of IRAK1^WT^ but not an IRAK1 variant lacking the target threonine, IRAK1^T387A^ (Kollewe et al., 2004), validating antibody specificity (Figure 3C-E and 3I, and Figure S3A). Importantly, kinase-dead (KD) and phosphomutant T209A IRAK1 variants (Kollewe et al., 2004) also failed to restore the anti-pT387 signals while phosphomimetic IRAK1^T209D^ enhanced immunoreactivity compared to WT (Figures 3F-H, 3I). These data indicated that IRAK1pT387 signals require both the activation and catalytic activity of IRAK1, and thus do indeed reflect autophosphorylation.

**Figure 3.**
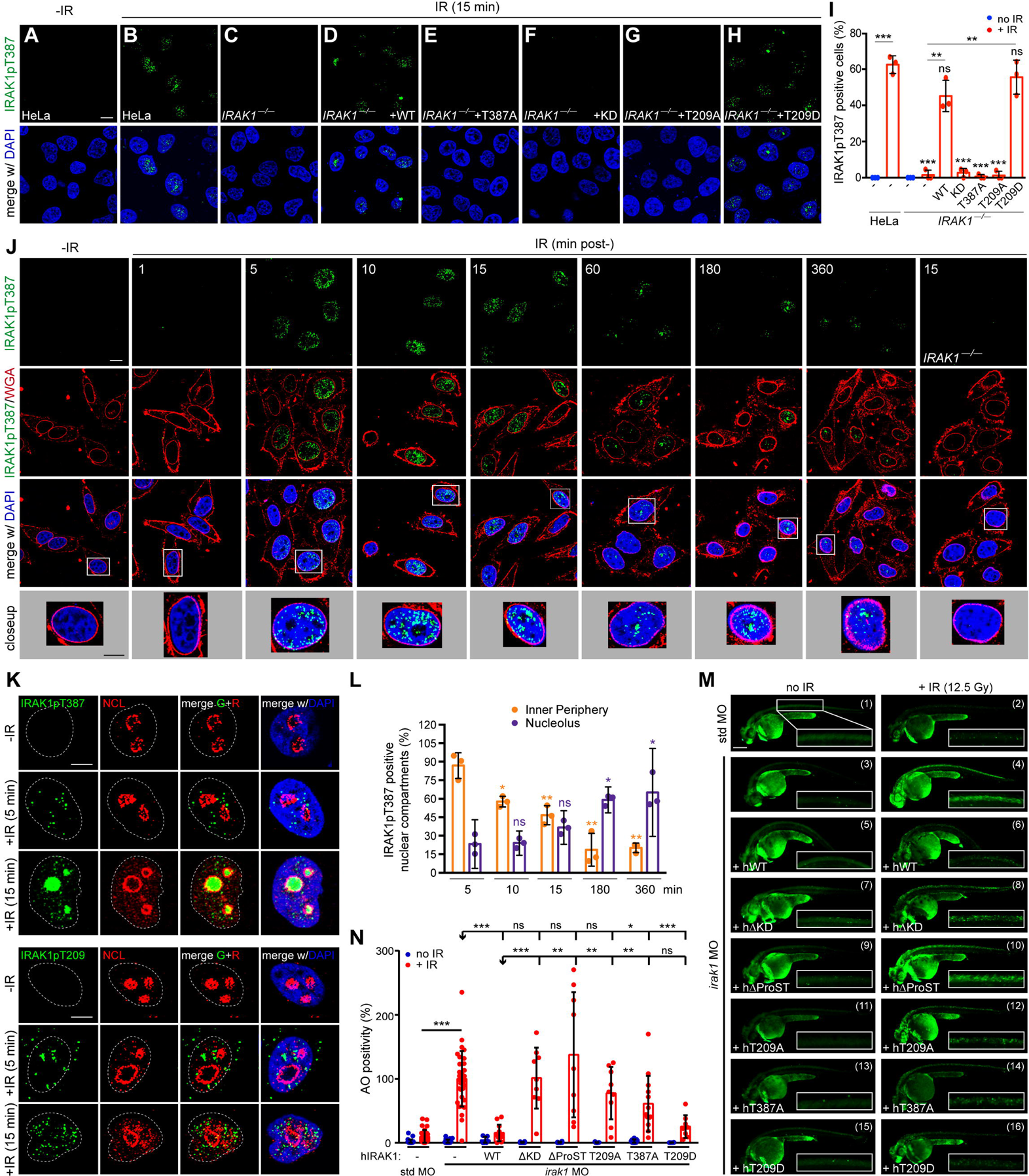
IRAK1 autophosphorylates in the nucleus of irradiated cells and relocates to nucleoli as a fully active kinase. (A-H) Parental and *IRAK1^—/—^* HeLa cells reconstituted with indicated FLAG-IRAK1 constructs were treated with or without IR (7.5 Gy), fixed 15 min after and stained with IRAK1pT387 antibody. Images representative of n=3 independent experiments were quantified in (I). Scale bar, 10 μm. (I) Quantification of IRAK1pT387 stains such as shown in (A-H). Data are means +/− SD of 3 independent experiments. \*\**p* < 0.005, \*\*\**p* < 0.001 ns, non-significant, two-tailed Student’s t-test. Expression levels of FLAG-IRAK1 variants shown in Figure S3B. (J) HeLa cells grown on cover slips were treated with or without IR (7.5 Gy), fixed at indicated time points (min) post-IR, stained with IRAK1pT387 antibody and co-stained with WGA and DAPI. Confocal images representative of *n*=3 independent experiments. Blowups of boxed areas indicate gradual relocalization of IRAK1pT387 to discrete nucleoplasmic areas. Scale bar, 10 μm. (K) HeLa cells grown on cover slips were treated with or without IR (7.5 Gy), fixed at indicated time points (min) after treatment, and stained with antibodies to nucleolin (NCL) and IRAK1pT387 (top panels) and IRAK1T209 (bottom panels). Images are close-ups of boxed areas from lower-magnification views showing multiple cells per condition (see Figure S3C-C’). Scale bar, 10 μm. (L) Quantification of images as shown in (J). Data are means +/− SD of 3 independent experiments. \*\**p* < 0.05, \*\**p* < 0.005, \*\*\**p* < 0.001 ns, non-significant, two-tailed Student’s t-test. (M) Zebrafish *p53^MK/MK^* embryos were co-injected at the 1 cell stage with the indicated MOs with or without synthetic mRNAs for the indicated human FLAG-IRAK1 variants. Embryos were treated with or without whole-body IR (12.5 Gy) 18 hours later and stained with acridine orange (AO) at 24 hpIR to label apoptotic cells in vivo. Spinal cord areas dorsal to the yolk tube (boxed in each panel) were used for quantification shown in (N). Scale bar, 0.2 mm. (N) Quantification of spinal cord areas (such as boxed in (M)) of at least 6 embryos per condition over at least 3 independent experiments. Data are means +/− SEM. Statistical significance vs. bars 2, 4 or 6: \**p* < 0.05, \*\**p* < 0.01, \*\*\**p* < 0.001, two-tailed Student’s t-test.

Examination of irradiated cells over time revealed discrete changes in the spatial distribution of IRAK1pT387 signals within the nucleoplasm (Figure 3J). The signal was first detected at the inner nuclear periphery (within 1 μm of the nuclear envelope), then spread throughout the nucleoplasm and eventually concentrated in discrete nuclear areas (Figure 3J). Co-staining with nucleolin and fibrillarin antibodies identified nucleoli as the destination of fully active IRAK1 (Figures 3K-L and S3C-D’). The localization of IRAK1pT387 to nucleoli reflected a specific process because partially active IRAK1 (IRAK1pT209) did not localize to that compartment and remained dispersed in the nucleoplasm (Figures 3K, bottom, and Figures S3C’,D’). The significance of the nucleolus to IR-induced IRAK1 signaling will be addressed below (see Figure 4K-N).

**Figure 4.**
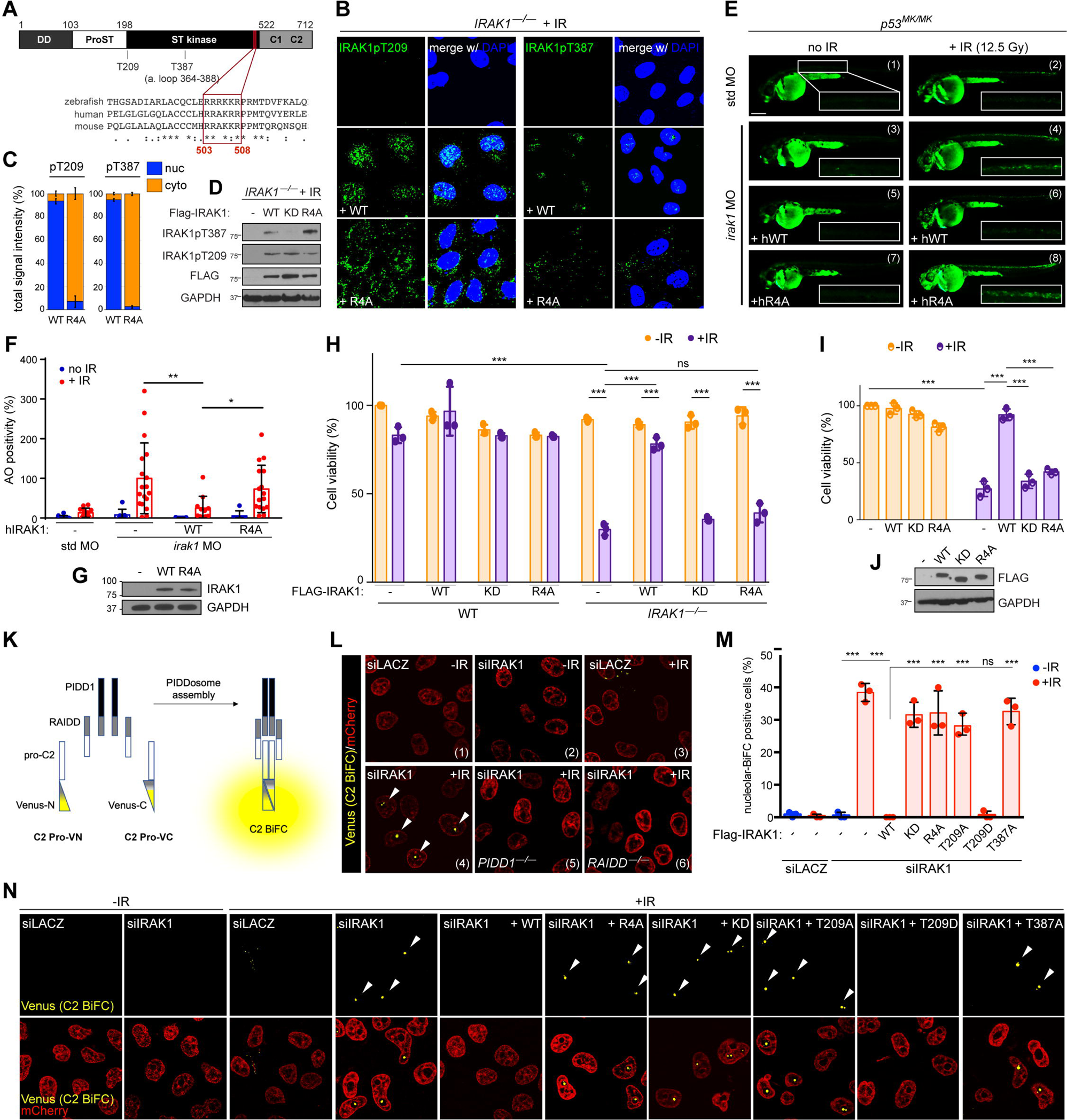
Nuclear internalization of active IRAK1 enables PIDDosome inhibition and is essential for IR-induced IRAK1 signaling. (A) Identification of a conserved putative NLS, R_503_RAKRR_508_, at the distal end of the IRAK1 kinase domain, with Clustal W alignment shown below. DD, death domain; ProST, proline/serine/threonine -rich domain; ST kinase, serine threonine kinase; C1 and C2, C-terminal regions 1 and 2. (B) *IRAK1^—/—^* HeLa cells reconstituted with indicated FLAG-IRAK1 constructs were treated with IR (7.5 Gy), fixed 15 min after, stained with IRAK1pT209 (left) or IRAK1pT387 (right) antibodies and counter stained with DAPI. Images representative of n=3 independent experiments were quantified in (C). R4A, mutation of all arginines in the putative human NLS shown in (A) to alanine (RRAKRR --+ AAAKAA). See also Figures S4C-D for expression levels of FLAG-IRAK1 variants and non-irradiated and parental controls. (C) Quantification of the spatial distribution of IRAK1pT209 (left) and IRAK1pT387 (right) signals from 3 independent experiments such as in (B). Blue, nuclear; orange, cytoplasmic. (D) Cells from (B) were lysed and analyzed by western blot (see also Figure S4C). (E) Zebrafish *p53^MK/MK^* embryos were co-injected at the 1 cell stage with the indicated MOs with or without synthetic mRNAs for the indicated human (h) FLAG-IRAK1 variants. Embryos were treated with or without whole-body IR (12.5 Gy) 18 hours later and stained with acridine orange (AO) at 24 hpIR to label apoptotic cells in vivo. Spinal cord areas dorsal to the yolk tube (boxed in each panel) were used for quantification shown in (F). Scale bar, 0.2 mm. (F) Quantification of spinal cord areas (such as boxed in (E)) of at least 6 embryos per condition over 3 independent experiments. Data are means +/− SEM, \**p* < 0.05, \*\**p* < 0.01, two-tailed Student’s t-test. (G) Pooled whole-embryo extracts from (E-F) analyzed by western blot to verify uniform expression levels of human FLAG-IRAK1 variants. The human IRAK1 antibody does not detect endogenous zebrafish Irak1. (H) Parental and *IRAK1^—/—^* HeLa cells reconstituted with indicated FLAG-IRAK1 constructs were treated with or without IR (7.5 Gy). Cells were stained with the vital dye alamarBlue 72 hr post-IR. Data are means +/− SD of 3 independent experiments. \*\*\**p* < 0.001; ns, non-significant; two-tailed Student’s t-test. Uniform expression of FLAG-IRAK1 constructs shown in Figure S4C. (I) Daoy cells, which are radiosensitive (compare bars 1 and 4), were reconstituted with indicated FLAG-IRAK1 constructs and treated with or without IR (7.5 Gy). Cells were stained with the vital dye alamarBlue 72 hr post-IR. Data are means +/− SD of 3 independent experiments. \*\*\**p* < 0.001; ns, non-significant; two-tailed Student’s t-test. (J) Cells from (I) were lysed and analyzed by western blot show uniform expression levels of FLAG-IRAK1 constructs. (K) Schematic of the caspase-2 bifluorescence complementation (C2 BiFC) assay. Cells expressing two fusion proteins containing the C2 prodomain (C2 Pro) tagged with either N- or C-terminal moieties of the Venus protein (C2 Pro-VN and C2 Pro-VC, respectively) emit the Venus signal when VN and VC are brought into close proximity as a result of dimerization of the C2 Pro domains. Such dimerization can result from PIDDosome complex assembly, in which case the release of the Venus signal genetically requires the scaffold protein PIDD1 and adaptor protein RAIDD, as is the case in (L). (L) C2.Pro-BiFC HeLa cells (Ando et al., 2017), which express C2 Pro-VN, C2 Pro-VC and nuclear mCherry, were transfected with indicated siRNA, treated with or without IR (10 Gy) and imaged 24 hours post-IR. Arrowheads mark Venus signal, reflecting dimerization of C2 Pro-VN and C2 Pro-VC. Note that depletion of IRAK1enables nucleolar C2 dimerization after IR (compare panels (3) and (4)) in a PIDD and RAIDD-dependent manner (compare (4), (5) and (6)). (M) C2.Pro-BiFC HeLa cells were transfected with indicated siRNA with or without indicated FLAG-IRAK1 variants, treated with or without IR (10 Gy) and scored for nucleolar C2 BiFC 24 hours post-IR. Data are means +/− SD of 3 independent experiments. \*\*\**p* < 0.001; ns, non-significant; two-tailed Student’s t-test. (N) Representative confocal images from (M). Arrowheads mark nucleolar C2 BiFC signals.

Similar to IRAK1^KD^ and IRAK1^T209A^, IRAK1^T387A^ failed to rescue *irak1* deficiency in irradiated *p53* mutant embryos (Figures 3M and 3N, compare bars 6 and 14; note that WT IRAK1 fully rescues Irak1-depleted fish, bars 4 vs. 6). Thus, IRAK1 autophosphorylation on T387 is necessary for IR-induced IRAK1 signaling. IRAK1^T387A^ did retain residual activity as compared with IRAK1^KD^ (Figure 3N, bars 4 vs. 15 (*p* < 0.05) and bars 4 vs. 8 (non-significant), respectively). This partial rescue indicated that while autophosphorylation on T387 is necessary for IR-induced IRAK1 signaling, it is unlikely the sole catalytic target of IRAK1 in the pathway.

### Nuclear translocation of active IRAK1 is essential for IR-induced IRAK1 signaling

Collectively, the data above supported a model for the early stages of IR-induced IRAK signaling where activation of IRAK4 in the cytoplasm leads to activation of IRAK1 therein, followed by its transport into—and full activation within—the nucleus. This was first supported by RNAi experiments which indicated that nuclear accumulation of IRAK1pT209 is sensitive to importin α/β dosage (Figure S4A,B). Nuclear translocation of active IRAK1 would be a defining feature of IR-induced IRAK1 signaling which, alongside un-reliance on MyD88, would distinguish the pathway from TLR/IL-1R signaling. To validate this notion, we tested whether nuclear transport even occurs, and if so, whether it is necessary IR-induced IRAK1 signaling.

Examination of mammalian IRAK1 sequences revealed a candidate nuclear localization sequence (NLS), RRAKRR, located at the distal end of the kinase domain which, notably, was conserved in zebrafish (Figure 4A). To disrupt the putative NLS, we mutated all arginines to alanine, thus generating an AAAKAA (R4A) IRAK1 variant. Introduction of FLAG-IRAK1^R4A^ into *IRAK1^—/—^* cells led to markedly reduced nuclear IRAK1pT209 and IRAK1pT387 signals after IR, with both the partially and fully activated species now trapped in the cytoplasm (Figures 4B-D and Figure S4D). Importantly, the R4A mutations did not compromise IRAK1 catalytic activity: IR-induced autophosphorylation on T387 was preserved in the variant, albeit occurring in the cytoplasm instead of the nucleus (Figures 4B-D and S4C,D). Thus, IRAK1^R4A^ represented a true separation-of-function mutant which uncoupled nuclear targeting from catalytic activity, disrupting the former while preserving the latter. This allowed us to specifically test whether IR- induced nuclear import of active IRAK1 is significant to IR-induced IRAK1 signaling.

In genetic complementation assays performed *in vitro* and *in vivo*, we found that IRAK1^R4A^ behaved as a function-null allele. Reconstitution of *IRAK1^—/—^* HeLa cells or Irak1- depleted zebrafish embryos with human IRAK1^R4A^ failed to rescue cell survival in response to IR, affording no added protection as compared to IRAK1^KD^ (Figure 4E-H). Furthermore, whereas WT IRAK1 was sufficient to protect radiosensitive Daoy cells from IR-induced cell death, IRAK1^R4A^ failed to do so, yet again mimicking IRAK1^KD^ (Figure 4I,J). Together, these experiments showed that nuclear translocation of active IRAK1 is essential for IR-induced IRAK1 signaling.

We next investigated the underlying mechanism. The proapoptotic PIDDosome complex (PIDD1-RAIDD-caspase-2) (Sladky et al., 2017; Tinel and Tschopp, 2004) is a downstream target of IR-induced IRAK1 signaling; PIDDosome inhibition by the kinase accounts, at least in part, for the pathway’s prosurvival function (Liu et al., 2019). Interestingly, PIDDosome assembly as triggered by DNA damage has been reported to occur primarily in nucleoli (Ando et al., 2017), that is, the destination of fully active IRAK1 in irradiated cells (see Figure 3 above).

We thus hypothesized that nuclear entry of active IRAK1 serves, at least in part, to enable PIDDosome inhibition by the kinase. To test this, we used the caspase-2 (C2) bimolecular fluorescence complementation (C2 BiFC) reporter system (Bouchier-Hayes et al., 2009), which probes PIDDosome formation in intact live cells (Figure 4K, see legend). As expected from previous studies (Liu et al., 2019), depletion of IRAK1 led to C2 BiFC in irradiated cells in a *PIDD1* and *RAIDD*-dependent manner (Figure 4L). While co-transfections with WT IRAK1 restored PIDDosome inhibition, IRAK1^R4A^ failed to do so (Figure 4M-N). IRAK1^R4A^ afforded no added protection as compared to IRAK1^KD^ or IRAK1 variants deprived of activation (T209A) or full activity (T387A) (Figures 4M-N). Thus, nuclear translocation of active IRAK1 is an essential feature of non-canonical IRAK1 signaling, critically required for the pathway’s anti-apoptotic function via PIDDosome inhibition.

### IR-induced IRAK1 signaling is triggered by DNA damage

Having identified a MyD88-independent, IRAK4—IRAK1 axis activated by IR, we sought to identify the upstream factors which initiate the pathway in place of the pathogen-sensing TLR/IL-1R–MyD88 module.

Despite the fact that all known TLR/IL-1R pathways which signal through IRAK1/4 do so via MyD88, we first addressed the possibility that a TLR/IL-1R might nevertheless be responsible for IR-induced IRAK1 activation. Indeed, pathways have been identified through which TLRs can signal independently of MyD88 (Fitzgerald and Kagan, 2020; O’Neill et al., 2013). While none so far have been shown to involve IRAKs, this could be the first instance of such a pathway. Secondly, irradiated cells may activate paracrine (and conceivably autocrine) TLR/IL-1R signaling via the release of (i) cytokines, PAMPs and DAMPs into the extracellular space which, in turn, might activate cell surface receptors; and, (ii) nucleic acids into the cytoplasm, which may activate intracellular TLRs (Apetoh et al., 2007; Candeias and Testard, 2015; Shan et al., 2007). However, medium transfer experiments in which non-irradiated cells were exposed to supernatant from irradiated donors ruled out an involvement of extracellular factors (Figure 5A-C, note that recipient cells fail to activate IRAK1). Similarly, depletions of all known intracellular TLRs—TLR3, TLR7, TLR8 and TLR9—failed to affect IR-induced IRAK1 activation (Figure S5A-D).

**Figure 5.**
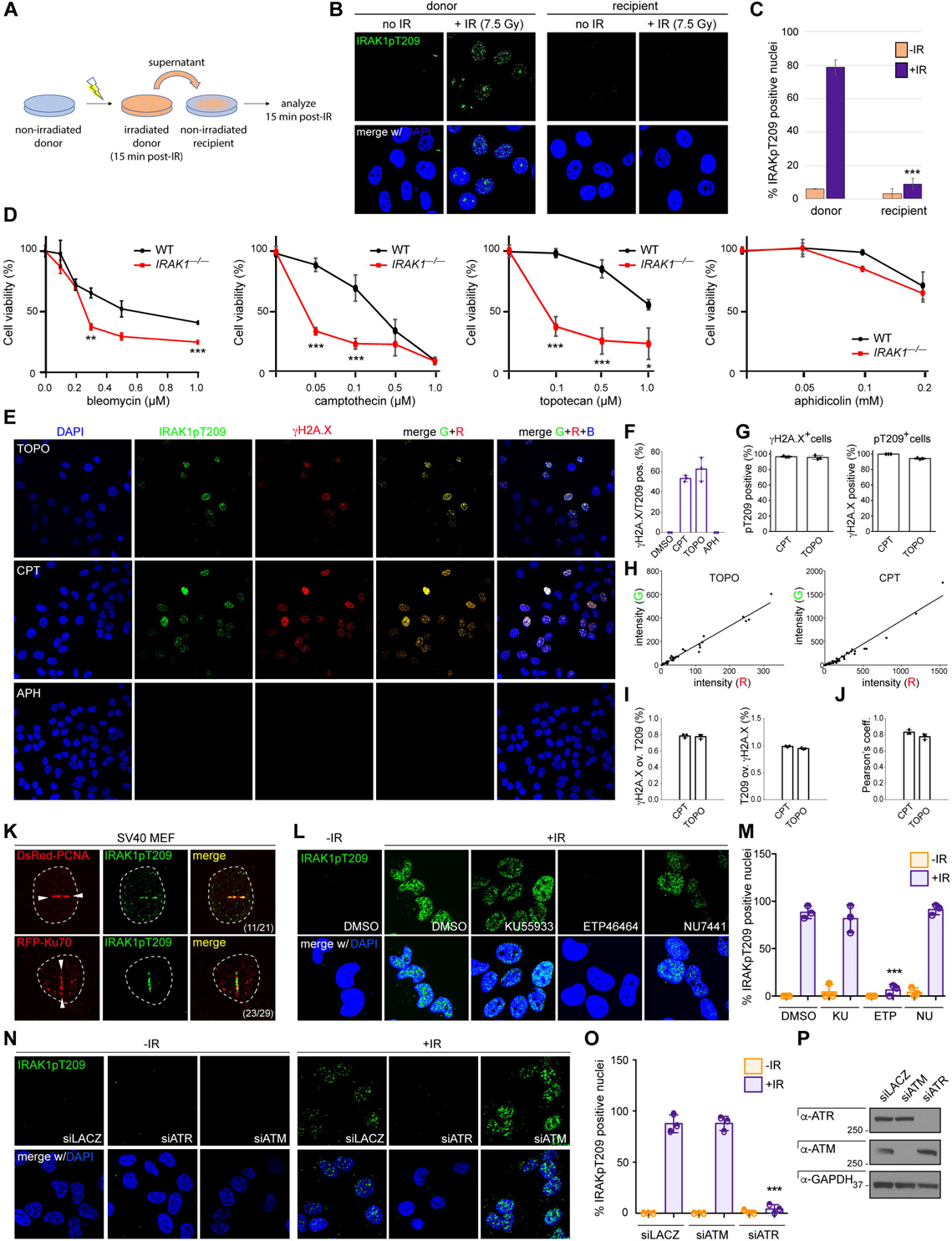
Non-canonical IRAK1 signaling is a DNA damage response. (A) Schematic of the medium-transfer experiments performed in (B-C). (B-C) HeLa cells treated as shown in (A) were stained with IRAK1pT209 antibody and DAPI, imaged by confocal microscopy (B), and data from 3 independent experiments were quantified (C). \*\*\**p* < 0.001; ns, non-significant; two-tailed Student’s t-test. (D) Parental and *IRAK1^—/—^* HeLa cells treated with indicated genotoxins at indicated doses were stained with the vital dye alamarBlue 72 hr post-treatment. Data are means +/− SD of 3 independent experiments. \*\*\**p* < 0.0001, two-tailed Student’s t-test. (E) HeLa cells treated with the indicated genotoxins were fixed after 6 hours and stained with indicated antibodies and DAPI. TOPO, topotecan (1 μM); CPT, camptothecin (1 μM); APH, aphidicolin (0.2 mM). Images representative of 3 independent experiments were analyzed in (F-J). See also Figure S5F-H for full time course and higher magnification views. (F) Cells double-positive for γH2A.X and IRAK1pT209 were counted from 3 independent experiments such as shown in (E). (G) The percentages of γH2A.X^+^ cells positive for IRAK1pT209 (left), and vice versa (right), were counted from 3 independent experiments as in (E). (H) Intensity of IRAK1pT7209 (G) signals relative to that of γH2A.X (R) in individual cells from 3 independent experiments as in (E) after treatment with TOPO (left) or CPT (right). (I) The percentages of γH2A.X^+^ foci positive for IRAK1pT209 (left), and vice versa (right), were counted from 3 independent experiments as in (E). (J) Pearson’s coefficient derived from image analyses in (I). (K) RFP-PCNA (top) and RFP-Ku80 (bottom) -expressing SV40 mouse embryonic fibroblasts (MEF) exposed to two photon laser micro-irradiation were stained with IRAK1pT209 antibody 15 min after treatment. DNA damage tracks indicated with arrowheads. Number of cells showing staining overlap between IRAK1pT209 and RFP-PCNA tracks (over 3 independent experiments) or RFP-Ku80 tracks or foci (over 2 independent experiments) are indicated. See Figure S5I for RFP-Ku80 foci. (L) HeLa cells exposed to the indicated DDR kinase inhibitors were treated with or without IR (7.5 Gy), fixed at 15 min post-IR and stained for IRAK1pT209 antibody and DAPI. Images representative of 3 independent experiments were analyzed in (M) . KU55933, ATMi (10 μM); ETP46464, ATRi (10 μM); NU7441, DNAPKi (5 μM). (M) Quantification of IRAK1pT209 stains from experiments as in (L). Data are means +/− SD of 3 independent experiments. \*\*\**p* < 0.001, two-tailed Student’s t-test. (N) HeLa cells transfected with indicated siRNA were treated with or without IR (7.5 Gy), fixed at 15 min post-IR and stained for IRAK1pT209 antibody and DAPI. Images representative of 3 independent experiments were analyzed in (O). (O) Quantification of IRAK1pT209 stains from experiments as in (N). Data are means +/− SD of 3 independent experiments. \*\*\**p* < 0.001, two-tailed Student’s t-test. (P) Whole cell lysates from cells as in (N) were analyzed by western blot with indicated antibodies.

In contrast, multiple independent lines of evidence pointed to IR-induced double strand DNA breaks (DSBs) as responsible for initiating the pathway. First, we had noted that the cytoplasmic activation and nuclear import of IRAK1 correlated with the presence of DNA damage in irradiated nuclei, with nuclear IRAK1pT209 signals declining coincident with DSB repair (12-24 hpIR) (Figure S2F-G). Second, drugs which, unlike IR, induce DSBs without causing the release of TLR agonists, such as the radiomimetic bleomycin and the topoisomerase inhibitors camptothecin and topotecan, were sufficient to trigger IRAK1 activation and its transport to the nucleus (Figures S5E-H, note that the DNA replication inhibitor aphidicolin had no effect). In turn, IRAK1 was required for the survival of the damaged cells, similar to its pro-survival role in irradiated cells (Figure 5D, again note that aphidicolin has no effect). Fourth, in these experiments, the numbers and intensity of nuclear IRAK1pT209 foci strictly correlated with the occurrence and extent of DNA damage, whether at a specific time point or as a function of time after stimulus (Figure 5E-H). In fact, a majority of nuclear IRAK1pT209 foci overlapped with sites of DNA damage (γH2A.X foci; Figures 5E, 5I-J and Figure S5F-H) or with DsRed-PCNA and RFP-Ku70 tracks or foci induced by 2-photon laser micro-irradiation (Figures 5K and S5I). Finally, interrogating the three major DSB-sensing kinases (ATM, ATR and DNAPK) (Blackford and Jackson, 2017) identified a requirement for ATR in IR-induced IRAK1 signaling. This was first seen with specific inhibitors, whereby an ATR inhibitor (ETP46464), but not ATM or DNAPKcs inhibitors (KU55933 and NU7441, respectively), blocked IR-induced IRAK1 activation (Figure 5L-M). These results were confirmed via RNAi silencing (Figure 5N-P). Collectively, these experiments identified DNA damage as the trigger of IR-induced IRAK1 signaling. Thus, the pathway differs from TLR/IL-1R signaling at the biological, molecular and subcellular levels, consistent with a non-canonical IRAK signaling pathway.

### E3 ubiquitin ligases Pellino1 and Pellino2 exert essential and distinct roles in IR-induced IRAK1 signaling

We next sought to gain first insight into the mechanism by which DNA damage induces IRAK4/1 activation independently of MyD88. The initiating step in the IRAK4-IRAK1 activation cascade, IRAK4 activation via *trans*-autophosphorylation (see Figure 1), requires proximity-induced dimerization (Cushing et al., 2014; Ferrao et al., 2014; Wang et al., 2017). Therefore, we reasoned that an adaptor or scaffold molecule would likely be required to enable IRAK4 dimerization in place of MyD88. A previously proposed substitute for MyD88, Unc5CL (Heinz et al., 2012), was dismissed because it showed no RNAi phenotype in human cells (Figure S6A-B) and is not conserved in zebrafish. We thus focused on other physical interactors of IRAK4 and IRAK1.

The E3 ubiquitin ligases Pellino1 (Peli1), Pellino2 (Peli2) and Pellino3 (Peli3) are homologs of *Drosophila* pellino, a protein originally identified as a physical interactor of the fly IRAK homolog, pelle (Grosshans et al., 1999). Human Peli proteins physically and functionally interact with IRAK1 *in vitro*, acting both as substrates of, and ubiquitin ligases for, IRAK1 (Moynagh, 2014; Zhang and Li, 2022). We focused on Peli1 and Peli2 because *PELI3^—/—^* cells could not be recovered despite multiple editing attempts.

*PELI1^—/—^* and *PELI2^—/—^* cells exhibited striking and distinct phenotypes. Ablation of Pel1 abrogated IR-induced IRAK1 activation altogether, whereas loss of Peli2 had no effect on activation but blocked the nuclear translocation of the activated kinase, trapping it in the cytoplasm (Figure 6A-B). In contrast, neither knockout had any discernable effect on IL-1β-induced IRAK1 activation (Figure 6A-B). The IR-specific phenotypes, which were confirmed by RNAi in multiple cell lines (Figure S6C), were rescued with respective GFP-tagged full-length proteins (Figure 6C). GFP-Peli1 failed to rescue *PELI2^—/—^* cells, and vice versa, confirming the non-redundant roles of either E3 ligase in IR-induced IRAK1 signaling (Figure 6C).

**Figure 6.**
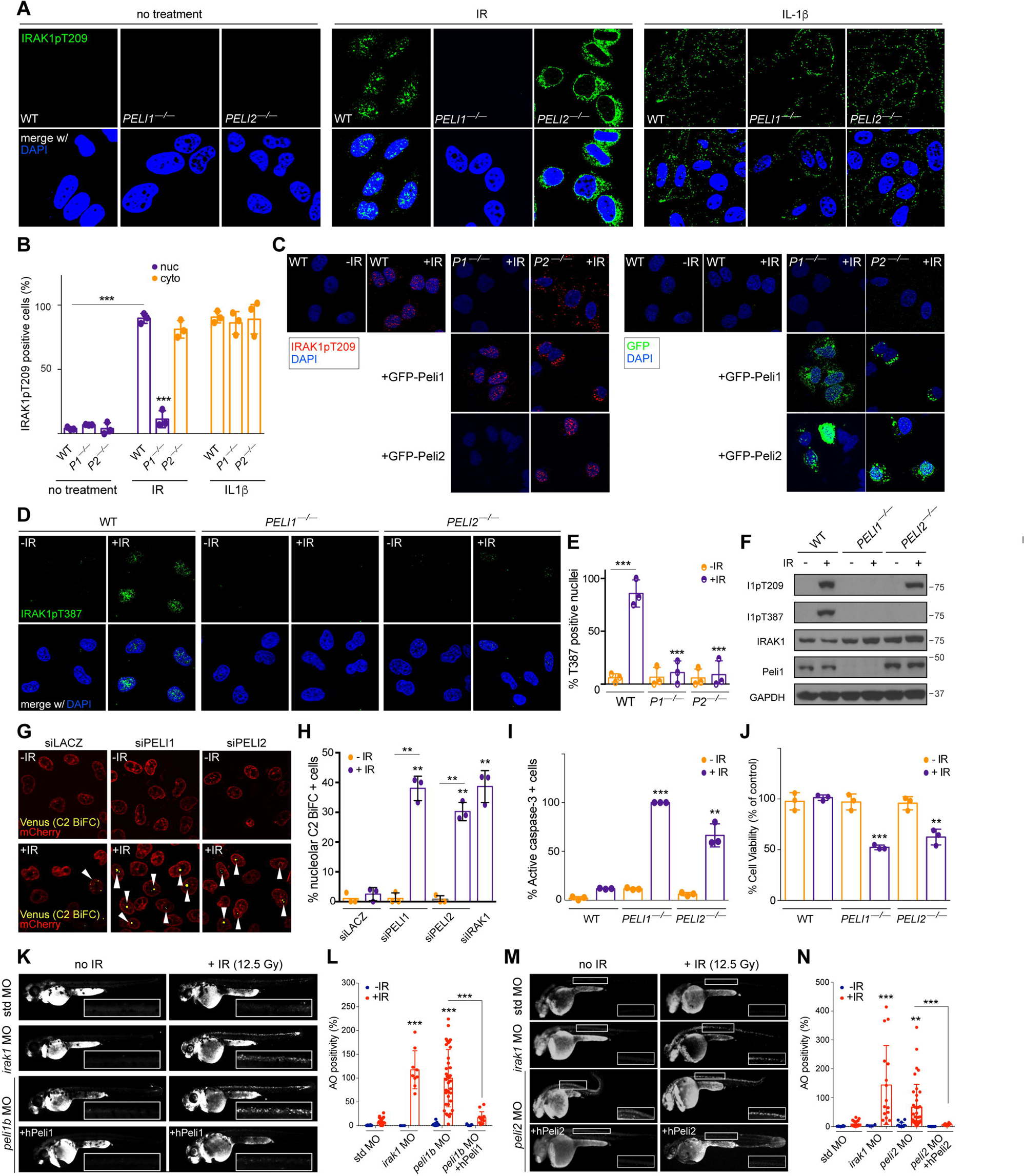
Essential and distinct roles of Peli1 and Peli2 in non-canonical IRAK1 signaling. (A) Parent *PELI1^—/—^* and *PELI2^—/—^* HeLa cells were treated with or without IR (7.5 Gy, middle) or IL-1β (0.1 μg/mL), right), fixed at 15 min post-treatment and stained with IRAK1pT209 antibody and DAPI. Images representative of 3 independent experiments were analyzed in (B). (B) Quantification of IRAK1pT209 signals in experiments as in (A). Purple, nuclear localization; orange, cytoplasmic localization. Data are means +/− SD of 3 independent experiments. \*\*\**p* < 0.001, two-tailed Student’s t-test. (C) Parent, *PELI1^—/—^* and *PELI2^—/—^* cells reconstituted with indicated GFP-Peli constructs were treated with or without IR (7.5 Gy), fixed 15 min post-IR, stained with IRAK1pT209 antibody (red) and DAPI, and additionally analyzed for GFP (green) by confocal microscopy. Images are representative of 3 independent experiments. (D) Parent, *PELI1^—/—^* and *PELI2^—/—^* cells were treated with or without IR (7.5 Gy), fixed at 15 min post-treatment and stained with IRAK1pT387 antibody and DAPI. Images representative of 3 independent experiments were analyzed in (E). (E) Quantification of IRAK1pT387 signals in experiments as in (D). Data are means +/− SD of 3 independent experiments. \*\*\**p* < 0.001, two-tailed Student’s t-test. (F) Whole-cell lysates from experiments as in (D) were analyzed by western blot with indicated antibodies. (G) C2.Pro-BiFC HeLa cells transfected with indicated siRNA were treated with or without IR (7.5 Gy) and fixed at 24 hours post-IR. Arrowheads mark nucleolar BiFC signals reflective of PIDDosome assembly (see Figure 4K-L). Images representative of 3 independent experiments were quantified in (H). (H) Quantification of BiFC signals in experiments as in (G). Data are means +/− SD of 3 independent experiments. \*\**p* < 0.005, two-tailed Student’s t-test. (I) Parent, *PELI1^—/—^* and *PELI2^—/—^* cells were treated with or without IR (7.5 Gy), fixed at 48 hours post-IR and stained with anti-active caspase-3 antibody and DAPI. Images representative of 3 independent experiments are shown in Figure S6D. Statistical significance vs. bar 2: \*\**p* < 0.005; \*\*\**p* < 0.001, two-tailed Student’s t-test. (J) Cells as in (I) were stained with the vital dye alamarBlue 72 hr post-IR. Data are means +/− SD of 3 independent experiments. Statistical significance vs. bar 2: \*\**p* < 0.005; \*\*\**p* < 0.001, two-tailed Student’s t-test. (K) Zebrafish *p53^MK/MK^* embryos were co-injected at the 1 cell stage with the indicated MOs with or without synthetic mRNA for human Peli1 (hPeli1). Embryos were treated with or without whole-body IR (12.5 Gy) 18 hours later and stained with acridine orange (AO) to label apoptotic cells in vivo. Spinal cord areas dorsal to the yolk tube (boxed in each panel) were used for quantification shown in (L). Scale bar, 0.2 mm. *peli1b*, zebrafish *PELI1* ortholog, as based on Clustal W phylogenetic analysis shown in Figure S6E. (L) Quantification of spinal cord areas (such as boxed in (K)) of at least 3 embryos per condition over at least 2 independent experiments. Data are means +/− SEM, \**p* < 0.05, \*\**p* < 0.01, \*\*\**p* < 0.001, two-tailed Student’s t-test. (M-N) As in (K-L) but substituting Peli2 for Peli1.

While Peli1 and Peli2 exerted distinct functions, loss of either ligase ultimately resulted in the same outcome: loss of nuclear accumulation of IRAK1pT209. Consistent with this, *PELI1^—/—^* and *PELI2^—/—^* cells both exhibited: (i) a complete loss of fully active IRAK1pT387 in irradiated nuclei (Figures 6D-F); and (ii) marked radiosensitive phenotypes, as evidenced by PIDDosome- mediated C2 activation (Figures 6G-H), downstream executioner caspase activation (Figures 6I and S6D), and reduced overall survival (Figure 6J) after IR. These phenotypes were recapitulated in *peli1b* and *peli2* morphant zebrafish (Figures 6K-N, note that each phenotype was rescued by co-injection with corresponding human WT mRNA; and Figures S6E-G). Thus, Peli1 and Peli2 are essential for IR-induced IRAK1 signaling, exerting non-redundant roles in the pathway.

### Pellino1 associates with IRAK4 and IRAK1 and acts non-catalytically to enable their activation after IR

Finally, we sought to investigate the mechanisms by which Peli1 and Peli2 promote the activation and nuclear transport of IRAK1, respectively. We focused on Peli1 because i) Peli1, but not Peli2, physically associated with IRAK1 after IR (Figure S7A-B); and, ii) none of the currently available anti-Peli2 antibodies specifically recognized Peli2, precluding biochemical studies at the endogenous level (see Methods). In contrast, the commonly used “Peli1/2” antibody specifically recognized Peli1 (Figure S7C).

To investigate Peli1, we first used a series of deletion- and catalytically-inactive GFP- Peli1 constructs (Figure 7A,B) (Ha et al., 2019). As expected from previous studies (Lin et al., 2008; Moynagh, 2014), GFP-Peli1 interacted with IRAK1 via the N-terminal forkhead-associated (FHA) domain (Figure S7D, note that GFP-Peli1^C^ is the sole construct that fails to interact with IRAK1). The FHA domain was essential for Peli1-mediated IRAK1 activation after IR (Figure 7C,D, note the lack of IRAK1pT209 signal in *PELI1^—/—^* reconstituted with GFP-Peli1^C^). In contrast, the catalytic RING-like domain was dispensable for Peli1 function (Figure 7C,D, note the rescue of *PELI1^—/—^* cells by GFP-Peli1^⊗C^). Likewise, each of three catalytically inactive Peli1 variants—H313A, H336A (Ha et al., 2019) and H369S/C371S (Schauvliege et al., 2006)—all restored IR-induced IRAK1 activation in *PELI1^—/—^* cells (Figure 7C,D). These observations indicated that Peli1 enables IR-induced IRAK1 activation through its interaction with, but not ubiquitination of, IRAK1.

**Figure 7.**
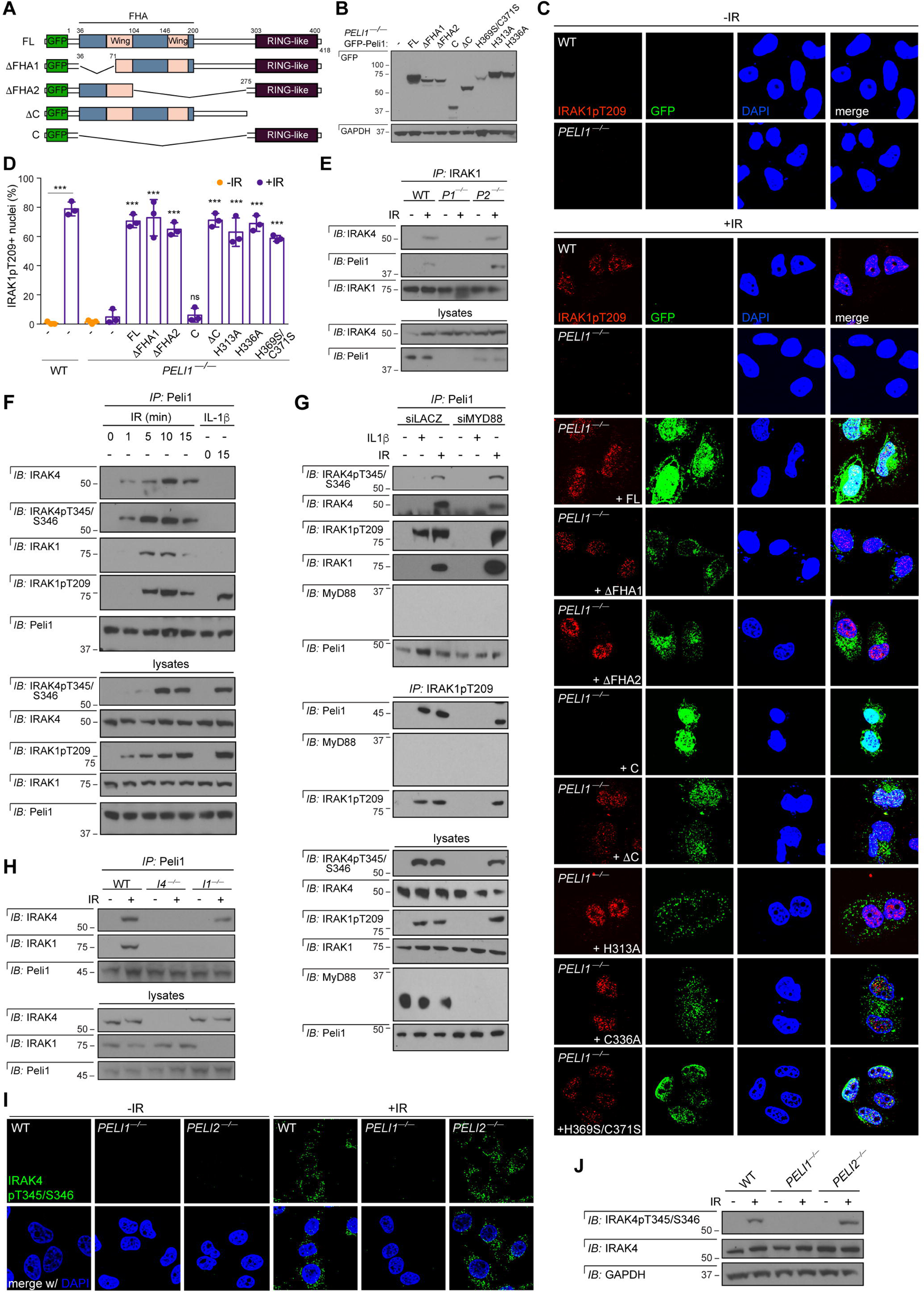
Peli1 functionally substitutes for MyD88 in non-canonical IRAK1 signaling. (A) Diagrams of full-length (FL) GFP-Peli1 and deletion constructs (Ha et al., 2019). FHA, forkhead-associated; RING-like, really interesting new gene-like (catalytic domain). ⊗FHA1 and ⊗FHA2 are partial deletions of the FHA domain and retain the ability to bind IRAK1, while the C construct is a complete deletion and fails to bind the kinase (see Figure S7D). ⊗C solely lacks the RING-like domain and binds IRAK1 (Figure S7D). (B) Expression levels of GFP-Peli1 constructs depicted in (A) in reconstituted *PELI1^—/—^* cells as analyzed in (C-D). Note that despite being expressed at relatively lower levels, the H369S/C371S mutant retains the ability to rescue *PELI1^—/—^* cells (see C-D). (C) Parent and *PELI1^—/—^* cells transfected with or without GFP-Peli1 constructs (see (A)) were treated with or without IR (7.5 Gy), fixed at 15 min post-IR, stained with IRAK1pT209 antibody (red) and DAPI, and imaged by confocal microscopy. GFP channel also shown. Images representative of 3 independent experiments were quantified in (D). (D) Quantification of IRAK1pT209 signals from (C). Data are means +/− SD of 3 independent experiments. Statistical significance vs. bar 4: \*\*\**p* < 0.001; ns, non-significant; two-tailed Student’s t-test. (E) Parent, *PELI1^—/—^* and *PELI2^—/—^* cells were treated with or without IR (7.5 Gy), harvested at 15 min post-IR, immunoprecipitated with IRAK1 antibody and analyzed by western blot with indicated antibodies. (F) Hela cells were treated with or without IR (7.5 Gy) or IL-1β (0.1 μg/mL), harvested at indicated time points, immunoprecipitated with Peli1 antibody and analyzed by western blot with indicated antibodies. (G) Hela cells transfected with indicated siRNA were treated with or without IR (7.5 Gy) or IL-1β (0.1 μg/mL), harvested at 15 min post-treatment, and Peli1 immunoprecipitates (top) and IRAK1pT209 immunoprecipitates (middle) were analyzed by western blot with indicated antibodies. (H) Parent, *IRAK4^—/—^* (*I4^—/—^*) and *IRAK1^—/—^* (*I1^—/—^*) cells were treated with or without IR (7.5 Gy), harvested at 15 min post-IR, and Peli1 immunoprecipitates were analyzed by western blot with indicated antibodies. (I) Parent, *PELI1^—/—^* and *PELI2^—/—^* cells were treated with or without IR (7.5 Gy), fixed at 15 min post-IR and stained with IRAK4pT345/S346 antibody and DAPI. Images representative of 2 independent experiments. (J) Cells as in (I) were harvested at 15 min post-IR and analyzed by western blot with indicated antibodies.

The structural role played by Peli1 suggested it might enable the IRAK4-IRAK1 interaction itself. Indeed, Peli1 was essential for this event whereas deletion of *PELI2* had no effect (Figure 7E). Peli1 associated with IRAK4 and IRAK1 as early as 1 min and 5 min post-IR, respectively (Figure 7F). This was in contrast to IL-1β-treated cells, in which Peli1 solely interacted with IRAK1 (active form) but not IRAK4, an observation in line with Peli1 serving as a substrate of IRAK1 in this context (Zhang and Li, 2022). IR-induced Peli1/IRAK4/IRAK1 complex formation did not require MyD88, in contrast to the IL-1β-induced Peli1-IRAK1pT209 interaction (Figure 7G). These results obtained through Peli1 pulldowns were confirmed in IRAK1pT209 immunoprecipitates (Figure 7G, middle blots). Thus, after IR, Peli1 associates with both IRAK4 and IRAK1 and functions non-catalytically to promote their interaction and subsequent IRAK1 activation.

These Peli1 properties were reminiscent of that of MyD88 in the TLR/IL-1R pathway, suggesting that Peli1 might act as the adaptor/scaffold (or subunit thereof) which substitutes for MyD88 in the DNA damage-induced pathway. In the immune pathway, MyD88 nucleates the MyDDosome by enabling the dimerization of IRAK4, which when autophosphorylated recruits, phosphorylates and activates IRAK1 (Wang et al., 2017). Therefore, should Peli1 function as the adaptor in the DSB-induced pathway, we would expect: (i) IRAK4 to be recruited to Peli1 prior to IRAK1; (ii) IRAK1 to be recruited to Peli1 via IRAK4; and (iii) Peli1 to be required for the activation of IRAK4 in addition to that of IRAK1. We found all three predictions to be true. First, Peli1 associated with IRAK4 as early as 1 min post-IR whereas IRAK1 was first detected in the complex at 5 min (Figure 7F). We also noted that IRAK4 autophosphorylation within the Peli1/IRAK complex initiates earlier than IRAK1 recruitment to the platform (Figure 7F). This observation was consistent with structural models implicating IRAK4 activation as a prerequisite for the IRAK4-IRAK1 interaction (Wang et al., 2017). Second, whereas IRAK1 was not required for the Peli1-IRAK4 interaction, IRAK4 was indispensable for IRAK1 recruitment to Peli1 (Figure 7H). Thus, Peli1 indeed associates with IRAK1 via IRAK4. Finally, IR-induced IRAK4 autophosphorylation on T345/S346 was completely dependent on Peli1 (Figure 7I), as further verified by western blot (Figure 7J). Collectively, these data identified Peli1 as a critical component of the MyD88-independent platform responsible for IRAK1/4 activation in irradiated cells (Figure S7E).

## Discussion

Here we delineate the backbone of a non-canonical pathway of IRAK kinase signaling distinct from the TLR/IL-1R pathway. While both pathways share a common mechanism for IRAK1 activation by IRAK4, non-canonical signaling differs both upstream and downstream of this core module (Figure S7E). DNA damage, not pathogen infection, triggers the pathway, via a Peli1- IRAK4-IRAK1 activation platform distinct from the MyDDosome. Once activated, IRAK1 does not remain in the cytoplasm but instead translocates to the nucleus where it functions, at least in part, to antagonize PIDDosome-mediated apoptosis. The pathway opens unforeseen areas of investigation in IRAK kinase signaling, DNA damage response (DDR) signaling and treatment resistance in cancer.

IRAK1 catalytic activity is largely dispensable for TLR/IL-1R signaling (Knop and Martin, 1999; Li et al., 1999; Maschera et al., 1999; Pauls et al., 2013), with the kinase instead acting structurally to activate TRAF6 and other potential effectors (Janssens and Beyaert, 2003). Yet in contrast with two of four IRAK family members, the pseudokinases IRAK2 and IRAK3, IRAK1 evolved to retain an intact catalytic domain (Lange et al., 2021; Muzio et al., 1997; Wesche et al., 1999). In the absence of a role for IRAK1 outside of innate immunity, the nature of the selective pressure to preserve its enzymatic activity remained elusive. Our discovery of a non-canonical pathway in which the kinase’s catalytic activity is essential may explain the evolutionary pressure to retain a functional catalytic site. We found that IRAK1 kinase function is, at minimum, required for autophosphorylation on T387, an event necessary for downstream anti-apoptotic signaling. Because IRAK1^T387A^ retained residual activity compared to a kinase dead variant, additional IRAK1 substrates are likely to be identified in the future.

Our study is the first to describe a physiological mechanism though which cells can activate IRAK4 and IRAK1 in a MyD88-independent manner. First evidence for the existence of such a mechanism came from a 2012 study by the Tschopp group which showed that overexpression of the DD protein UNC5CL could activate NF-κB and JNK signaling in an IRAK1- and IRAK4-dependent but MyD88-independent manner (Heinz et al., 2012). Because UNC5CL physically associated with IRAK4 and IRAK1, the group proposed that UNC5CL might provide an alternate scaffold for IRAK4 and IRAK1 activation. However, these observations still await genetic and physiologic validation, and we have ruled out a role for UNC5CL in non-canonical IRAK signaling (Figure S6A,B). Instead, we identified a non-DD-containing but extensively validated interactor of IRAK kinases, Peli1, as the likely adaptor protein substituting for MyD88. This was surprising given that Peli proteins are commonly viewed as downstream effectors of IRAK1, acting both as substrates of, and E3 ubiquitin ligases for, the active kinase (reviewed in Moynagh, 2009; Schauvliege et al., 2006; Zhang and Li, 2022). In line with such a downstream role in canonical IRAK signaling but an upstream role in non-canonical signaling, removal of Peli1 or Peli2 had no effect on IL-1β-induced IRAK1 activation in experiments where irradiated *PELI1^—/—^* and *PELI2^—/—^* cells analyzed in parallel showed profound phenotypes. Furthermore, the E3 ligase activity of Peli1 was irrelevant to its function in IRAK1/4 activation, and evidence of polyubiquitination of either kinase was lacking in all IRAK1/4 pulldowns and lysates analyzed in this study. The non-catalytic role played by Peli1 echoes earlier studies in the Pellino field where Peli1-3 were primarily viewed as scaffolding proteins orchestrating protein interactions in the TLR/IL-1R pathway (Schauvliege et al., 2007). Our data are consistent with a model where Peli1 first associates with IRAK4 to promote its activation, followed by IRAK1 recruitment to the Peli1/IRAK4 complex and its activation by IRAK4 therein. As such, the Peli1/IRAK4/IRAK1 complex would mirror the activity of the MyDDosome (Wang et al., 2017; Wang et al., 2006). Future studies will determine whether Peli1, like Myd88, acts by enabling the dimerization of IRAK4, whether Peli1 acts a subunit of a larger complex in which another molecule is responsible for dimerization, or whether the Peli1-IRAK4 interaction stimulates IRAK4 activation via an as-yet unidentified mechanism.

Interestingly, Peli1 has been implicated in the DDR and, notably, appears to localize to DSBs in response to IR (Dai et al., 2019; Ha et al., 2019), where it undergoes ATM-mediated phosphorylation within minutes of treatment (Ha et al., 2019). In turn, phosphorylated Peli1 ubiquitinates NBS1 to regulate DSB repair via homologous recombination. We found that IRAK1pT209 also localizes at DSBs in response to IR with similar timing, which suggested that DSBs might be the site for Peli1-mediated IRAK1 activation in non-canonical signaling. However, this was ruled out by multiple lines of evidence identifying the cytoplasm as the site of IRAK1 activation in irradiated cells. We also considered the hypothesis that a fraction of the ATM- phosphorylated Peli1 protein pool (Ha et al., 2019) might vacate the nucleus to direct IRAK4/IRAK1 activation in the cytoplasm, thus providing a mechanism by which DSBs instruct Peli1/IRAK4/IRAK1 complex formation in that compartment. However, we could not detect any changes in the spatial distribution of endogenous Peli1 within the time window of IR-induced IRAK1 activation (0-15 min post-IR), with the protein remaining diffusely expressed throughout the nucleus and cytoplasm (our unpublished observations). Additionally, ATR, not ATM, is the DDR kinase responsible for engaging non-canonical IRAK signaling. Despite these discrepancies, our study adds to that of others to further implicate Peli1 as a DDR effector (Dai et al., 2019; Ha et al., 2019). How DSBs and ATR in the nucleus ultimately direct Peli1/IRAK4/IRAK1 platform assembly in the cytoplasm, whether through Peli1 itself or another mechanism, is the next fundamental question in the field. Additionally, while we could tie the nucleolar localization of fully active IRAK1 to its action as a PIDDosome inhibitor, the significance of the localization of IRAK1pT209 at DSBs remains to be explored. Given the pro-survival role of the pathway, a function in DSB repair seems plausible.

Another outstanding question is the mechanism by which Peli2 enables the transport of activated IRAK1 to the nucleus. In response to DNA damage, hundreds of proteins are promptly transported to the nucleoplasm where they perform essential functions as DNA repair effectors, transcription factors or replication regulators (Bennetzen et al., 2018; Fabbro and Henderson, 2003; Knudsen et al., 2009). However, strikingly little is understood about the underlying mechanisms: How the occurrence of DNA injury in the nucleus is conveyed to or sensed in the cytoplasm and, in turn, how the nuclear import machinery recruits target cargos for transport, all within minutes of stimulus, remain largely unexplored. This applies to major tumor suppressors such as p53 and PTEN, whose nuclear internalizations are nevertheless essential for tumor suppression (Marine, 2010; Trotman et al., 2007). The Peli2 null phenotype provides a new entry point into this biology, whereby the nuclear transport of IRAK1pT209 is completely and specifically blocked upon loss of a single E3 ligase. Because ubiquitination is critical to the import of PTEN (Trotman et al., 2007) and p53 (Marchenko et al., 2010; Nikolaev et al., 2003; Yamasaki et al., 2007), it is tempting to speculate that Peli2-mediated ubiquitination of IRAK1 might promote its IR-induced nuclear translocation. While Peli2-mediated ubiquitination of IRAK1 has been extensively described in other settings (reviewed in (Moynagh, 2014)), we have not thus far detected an interaction between Peli2 and IRAK1 in response to IR (Figure S7A,B). To the best of our knowledge, antibodies which specifically recognize endogenous Peli2 have not been identified, including by us over the course of this study, but will be required to address these questions.

Our previous work implicated IR-induced IRAK1 signaling as a driver of tumor-intrinsic R-RT, thus identifying IRAK1 as a therapeutic target in radiation-resistant cancers (Liu et al., 2019; Liu and Sidi, 2019). IRAK1 inhibitors showed efficacy in zebrafish and mammalian models of tumor R-RT at doses tolerated by non-irradiated tissues. The therapeutic window suggested by this data, in line with the absence of overt phenotypes in *Irak1^—/—^* mice (Thomas et al., 1999), overcomes the limitations of traditional radiosensitizers and is of urgent need in the clinic (Lawrence et al., 2013; Sharma et al., 2016). Our present work identifies multiple immunofluorescence markers of RT-induced IRAK1 signaling which could be used as diagnostic tools to predict IRAK1i treatment efficacy in patient biopsies, as well as new pathway members (IRAK4, Peli1, Peli2, ATR) which might define additional targets for inhibition. Finally, our work reveals that tumor cells can engage non-canonical IRAK1 signaling not only in response to IR- induced DNA damage but also upon exposure to DSB-inducing genotoxins such as radiomimetics and topoisomerase inhibitors. IRAK1 was notably essential for the survival of cancer cells treated with such drugs, suggesting a broad applicability of IRAK1 inhibitors in treatment-resistant cancers.

## Supporting information

Li Shah et al Supplement

## Acknowledgements

We thank Carlos Franco and Diarles Carles for zebrafish care, and Michael Cohen, Vicky Rao, Xiaoxia Li, Jonathan Ashwell and Chang-Woo Lee for critical reagents. Y.L. was supported by a National Cancer Center postdoctoral fellowship award. S.Z. was supported by NIH/NCI RO1CA275184. S.S. was supported by grants from NIH/NIGMS (RO1GM135301), NIH/NCI (R01CA178162), Searle Scholars Program, Pershing Square Sohn Cancer Research Alliance and New York Community Trust.

## Author contributions

Y.L. and R.B.S. performed most zebrafish and cell culture experiments, respectively, with contributions from S. Sarti. (IRAK4pT345/S346 time course, Peli2 MO and importin studies), A.L.B. (IRAK1pT387 time course), A.G. and J.M.S. (CRISPR- Cas9 editing), F.N. (assistance with IRAK4pT345/S346 IF), I.Y. (assistance with clonogenic assays), Z.S. (assistance with micro-IR image analysis), and B.J.L, Z. Shao and S.Z (2-photon micro-IR treatments). S.S. designed and supervised the study, and wrote the manuscript with Y.L., R.B.S and S. Sarti. All authors discussed the results and participated in the manuscript preparation and editing.

