## Supplementary material for "A Non-Canonical IRAK Signaling Pathway Triggered by DNA Damage": Li Shah et al Supplement

### Supplementary Information

Li, Shah et al.

**Materials and Methods**

**Supplementary Legends**

**Supplementary References**

**Supplemental Table 1**

**Supplementary Figures 1-7**

#### Materials and Methods

##### Cell Culture, Reagents and X-Ray Irradiation

HeLa (cervical, p53-degraded via HPV-E6), CAL27 (HNSCC, TP53 H183L), SAS (HNSCC, TP53 E336*), MCF7 and MDA-MB-231 (breast, TP53 R280K) cell lines were cultured in DMEM medium (Life Technologies) supplemented with 10% Fetal Bovine Serum (FBS) (Sigma-Aldrich) and 1% penicillin–streptomycin (P/S) (Life Technologies). Daoy (radiosensitive MB cell line) cells were cultured in EMEM medium (Life Technologies) supplemented with 10% FBS and 1% P/S. HeLa, CAL27, Daoy cells were obtained from American Type Culture Collection. SAS cells were purchased from Health Science Research Resources Bank. MDA-MB-231 and MCF7 cells were provided by D. Germain. C2.Pro-BiFC cells (Ando et. al. JCB, 2017) were cultured in DMEM medium (Life Technologies) supplemented with 10% FBS and 1% P/S. ETP46464, camptothecin, aphidicolin and bleomycin were purchased from Sigma-Aldrich. KU-55933 was purchased from TOCRIS. Topotecan and Nu7441 were purchased from Selleckchem. Recombinant human IL-1β was from PeproTech (200-01B). For X-ray IR, cells seeded on 10 cm, 6 well or 96-well plates were irradiated at the indicated doses using the Precision X-rad 320 irradiator. The plates were placed on the programmable shelf with the turntable left on. Filter 1 (2mm Al) and Program 5 (platform height of 50 cm from the source, with an offset of 6 cm and a dose rate of 320 kV) were used in all experiments.

##### CRISPR–Cas9 Gene Editing

Plasmid lentiCRISPR v2 was digested with BsmBI-v2 (New England Biolabs, R0580) according to the manufacturer's recommendations. Briefly, 1 µg of plasmid was digested for 1 h at 55 °C, then digested plasmid was gel purified using a QIAquick Gel Extraction kit and eluted in water. Single guide RNA (sgRNA) oligonucleotides for cloning were annealed by mixing them in equal 10 µM concentrations with the addition of 1×T4 DNA ligase buffer, then the mixture was incubated at 37 °C for 30 min, then at 95 °C for 5 min and then ramped down to 25 °C at 5 °C $min^{-1}$. Hybridized oligonucleotides were diluted 1:200 with $H_2O$. BsmBI-v2 digested lentiCRISPR v2 plasmid (50 ng) was ligated with 1 µM final concentration oligo duplex using T4 Ligase (NEB, M0202) according to the manufacturer's recommendations and incubated overnight at 4 °C. XL-1 blue competent *E. coli* were transformed with 1 µl of ligation reaction according to the manufacturer's protocol (Agilent Technologies, catalogue no. 200249). Single clones were sequence verified using Sanger sequencing. Lentivirus particles containing sgRNA constructs of *IRAK4*, *PELI1*, *PELI2* and *PELI3* were generated by transfecting Phoenix packaging cells with lentiCRISPR v2 containing corresponding sgRNAs (Eurofins Genomics) and a combination of the lentiviral helper plasmids pCMV-dR8.91 and pMD.G at a ratio of 2:1:1, respectively. jetPEI (Polyplus, 101-10N) was used as the transfection ant. After 24 h, medium containing viral particles was collected and concentrated using Lenti-X Concentrator according to the manufacturer's protocol (Clontech, 631231). Briefly, 1 volume of Lenti-X Concentrator was mixed with 3 volumes of 0.45-µm filtered viral particle-containing media. The solution was then incubated overnight at 4 °C. The samples were centrifuged at 1,500 × *g* for 45 min at 4 °C, supernatant aspirated and the pellet resuspended in HeLa cell culture media. For infection, $2 \times 10^5$ HeLa cells were plated into 6-well plates. The next day, 5 µM polybrene (Millipore, tr-1003-g) and 200 µl of concentrated viral particles were added per well. The medium was replaced the next day with medium containing 1 µg $ml^{-1}$ puromycin for selection. *IRAK1*$^{-/-}$ HeLa cells were generated and validated previously (Liu et al., 2019). *PELI3*$^{-/-}$ could not be recovered. sgRNA sequences are listed in Supplementary Table 1.

**RNAi**

siRNA transfections were performed using X-tremeGENE siRNA transfection reagent (Roche) and 20 nM siRNA according to the manufacturer's instructions. Cells were treated with IR (7.5

Gy) or IL1β (0.1 μg/ml) at 48 hrs post-transfection. Cells were harvested for either western blotting, co-immunoprecipitation or Immunofluorescence 15 mins post-treatment unless otherwise described. Previously validated siRNAs were siLACZ (Sidi et al., 2008), siIRAK1, siIRAK4, siMYD88 (Liu et al., 2019), siATM and siATR (Ando et al., 2012)(Qiagen). siRNAs targeting Pellino1, Pellino2, UNC5CL, TLR3, TLR4, TLR7, TLR8, TLR9, Importin-α2 and Importin-β1 were purchased from Qiagen. siRNA sequences are listed in Supplementary Table 1.

**Plasmids and DNA Transfection into Cultured Mammalian Cells**

pVQAd-IRAK4 WT, D329A and R12C, and pcDNA3-flag-IRAK1 WT, T387A, KD, T209A, T209D, DKD and DProST, were generous gifts from Vicky Rao (De et al., 2018), Michael Martin (Kollewe et al., 2004), Xiaoxia Li (Yao et al., 2010) and Jonathan Ashwell (Conze et al., 2008). IRAK1$^{R4A}$ was generated from pcDNA3-flag-IRAK1$^{WT}$ using a Q5® Site-Directed Mutagenesis Kit (NEB, E0554S) with the primers IRAK1_R4AF 5'-aaagcagcaCCTCCTATGACCCAGGTGTACG-3' and IRAK1_R4AR 5'-ggccgcggcGTGCAGGCAGCAGCAGGC-3', resulting in R$_{503}$RAKRR$_{508}$ mutated to A$_{503}$AAKAA$_{508}$. pEGFP-C1-Flag-Peli1 and derived deletion constructs DFHA1, DFHA2, DC, C, and catalytically inactive mutants H313A and H336A were generous gifts from Chang-Woo Lee (Ha et al., 2019). The H369S/C371S double-mutant, homologous to the catalytically dead Peli2 H371S/C373S variant (Ordureau et al., 2008), was generated from pEGFP-C1-Flag-Peli1 by site-directed mutagenesis using primers Peli1-H369C371S-Q5Fw 5'-GAGCTCAGAAAAGACAACTGCCTATTGGTC-3' and Peli1-H369C371S-Q5Rv 5'-ACGCTCCCACACGGGCTAAACGC-3'. Full-length *PELI2* cDNA (NM_021255.3) was amplified by 2 rounds of PCR using Phusion Flash High-Fidelity PCR Master Mix (Thermo Scientific). The first round was performed using a primer pair within the flanking UTRs of *PELI2*, and the second round was a NESTED PCR. Primers were Peli2-5'UTR-Fw 5’-GCTGCTGTTTTGAGCATGCAA-3’, Peli2-3'UTR-Rev 5’-ACTGGCCGCCCCGCGCCCCCTT-3’, Peli2-XhoI-Fw 5’-ATACTCGAGGCCACCATGTTTTCCCCTGG-3’ and Peli2-HindIII-Rev 5’-AAGCTTTCAGTCAATTGGACCTTGGA-3’. Full-length *PELI2* cDNA was cloned in the pGem®-T easy (Promega) according to the manufacturer’s instructions, and was subcloned downstream of EGFP into XhoI and HindIII sites of pEGFP-C1-Flag to obtain pEGFP-C1-Flag-Peli2. All primers for site-directed mutagenesis were designed using NEBaseChanger (https://nebasechanger.neb.com) and all targeted mutations were verified by DNA sequencing. Plasmid DNA transfections were performed using X-tremeGENE HP DNA Transfection Reagent (Roche) with a ratio 2:1 according to the manufacturer’s instructions. With the exception of pEGFP-C1-Flag-Peli1$^{FL}$ in Figure 7B-D (0.5 μg/mL), all plasmids were transfected at 1 μg/mL. pDsRed-PCNA (derived from pDsRed, Clontech) and pRFP-Ku70 (derived from pRFP-C/N serial, Clontech) expression plasmids were used in two-photon micro-IR experiments (see below) and were generously provided by Dr. David Chen from the University of Texas Southwestern and Dr. Li Lan from Massachusetts General Hospital, respectively.

**Cell Viability Assays**

AlamarBlue-based cell viability assays were performed as described (Liu et al., 2019) with several modifications. Cells were seeded into 96-well plates at a density of 400 cells/well. After 16 hours, cells were treated with bleomycin, camptothecin, topotecan and aphidicolin at their indicated doses or 7.5 Gy IR. 3 days post-treatment, cells were incubated with alamarBlue (Thermo Fisher) at a final concentration of 10%. Absorbance was measured at a wavelength of 570 nm with a 600 nm reference wavelength. Relative fluorescence (RFU) was calculated using cell free wells as a control reference and percent survival was calculated compared to DMSOtreated, non-treated controls. Reverse transfections were performed for siRNAs while seeding the cells. cDNA transfections were performed 24 hrs prior to IR treatment.

**Clonogenic Assays**

Single-cell suspensions were seeded into 6-well plates (50-200 cells/well) and irradiated at indicated doses. After being cultured for 12 days, plates were rinsed with PBS, incubated with fixing solution (75% methanol, 25% acetic acid) and stained by 0.5% crystal violet (Sigma-Aldrich, St Louis, MO, USA) in methanol for 30 min at room temperature. Colonies consisting of at least 50 cells were scored.

**Immunofluorescence and Confocal Microscopy (Tissue Culture)**

Cells ($1 \times 10^5$) were seeded on coverslips in 6-well plates, fixed in 1% paraformaldehyde, permeabilized in 0.25% Triton X-100, blocked in 1% BSA–PBS for 30 mins at 37$^{o}$c, stained with indicated primary antibodies (for dilutions, see Supplementary Table 1) for 2 hours at 37 $^{o}$C and secondary antibody (1:200, anti-rabbit, AlexaFluor 488, AlexaFluor 555, AlexaFluor 647 Invitrogen), mounted in Vectashield with 4,6-diamidino-2-phenylindole (DAPI) and sealed with nail polish, as described previously. Images were obtained under a ×63 NA 1.40 oil objective with an inverted confocal microscope (405 nm, 488 nm, 647 nm; SP5, Leica) and acquired using LAS software. For experiments involving WGA staining, cells were stained with WGA at a dilution of 2.5:1000 prior to permeabilization.

**Image Analysis**

Cytoplasmic/nuclear Intensity quantification was performed using Intensity Ratio Nuclei Cytoplasm Tool (RRID:SCR_018573) URL:

https://github.com/MontpellierRessourcesImagerie/imagej_macros_and_scripts/wiki/Intensity-Ratio-Nuclei-Cytoplasm-Tool, installed as a plug-in in FIJI software. Co-localization analyses were performed with Just Another Co-localization Plugin (JaCoP) https://imagej.net/plugins/jacop, downloaded as a plugin for FIJI. Pearson's coefficient was calculated as a co-localization indicator.

**Western Blotting and Antibodies**

Cells seeded at a density of 0.5 $x10^6$ in 10 cm plates were grown to 60–70% confluence. Cells were transfected with indicated siRNAs for 48 hours and cDNAs for 24 hours. For experiments involving Irradiation, cells were treated with 7.5 Gy IR for 15 minutes post-transfection, harvested and lysed using 1% NP-40 buffer (Boston BioProducts) with protease and phosphatase inhibitors. Lysates (50 – 200 μg) were incubated at 70°C for 10 minutes after adding NuPAGE LDS Sample Buffer (4X) (Life Technologies) and 5% 2-Mercaptoethanol (Sigma Aldrich). Samples were run on a 10% Bis-tris gel (Nupage) at 170V for 1 hour. After electrophoresis, samples were transferred on a nitro-cellulose membrane (Thermo Fisher Scientific) at 94 V for 100 minutes. Membranes were then blocked with 5% Bovine serum albumin (BSA, Sigma Aldrich) in TBS with 0.1% Tween and probed with primary antibodies at 4°C overnight. Membranes were then rinsed with TBS-Tween (5x5 min) and probed with specific HRP-linked secondary antibody in 5% milk or BSA (in TBS-Tween) for 1 hr at room temperature. Membranes were washed as described earlier and placed in SuperSignal West Pico Chemiluminescent Substrate or SuperSignal West Dura Extended Duration Substrate (Pierce Biotechnology). The membrane was then developed with photographic film. A full list of antibodies and protocols can be found in Supplementary Table 1. The "Peli1/2" (clone F-7) commercialized by Santa Cruz Biotechnology and other companies recognizes Peli1, but not Peli2 (Figure S7C). To our knowledge, no commercial or custom antibody has been reported that specifically detect Peli2, and none of the three antibodies commercialized as specific to

Peli2 (see Supplementary Table 1) recognized Peli2 when tested as in Figure S7C (our unpublished observations).

**Co-Immunoprecipitation**

Lysates for immunoprecipitation (IP) were prepared in 1 or 0.1% NP-40 buffer (1 or 0.1% NP-40, 50 mM Tris-HCl [pH 8.0], 150–250 mM NaCl, 5mM EDTA, 1 mM phenylmethylsulfonyl fluoride, protease inhibitors cocktail [Complete Mini, Roche] and phosphatase inhibitor cocktail [PhosSTOP, Roche]). For endogenous IPs, whole-cell lysates (1–5 mg) were mixed with Protein-G magnetic beads (Invitrogen, 20 μl of a 50% slurry) and IRAK1 and Pellino1 (5 mg, 1 ml final volume) for 10 min to 1 hour at room temp on a rotating wheel. Beads were then washed three times with PBS-Tween20 (0.02%), resolved by SDS-PAGE, and probed with primary antibodies detected with the corresponding secondary antibodies or mouse TrueBlot HRP-conjugated secondary antibodies [eBioscience]). For α-Flag or α-GFP IPs, whole-cell lysates (0.15–2 mg) were mixed with 20 μL beads (50% slurry) and mouse α-Flag (M2) antibody (3 mg) or α–GFP (3 mg) in 1% NP-40 buffer (500 mL final volume) for 10 min on a rotating wheel. Beads were then washed three times with PBS-T, resolved by SDS-PAGE and analyzed by western blot.

**Caspase-2 Bimolecular Fluorescence Complementation (C2 BiFC) Imaging**

Bimolecular fluorescence complementation (BiFC) uses nonfluorescent N- and C-terminal fragments of the yellow fluorescent protein Venus ("split Venus") that can associate to reform the fluorescent complex when fused to interacting proteins (Shyu et al., 2006). When the C2 prodomain is fused to each half of split Venus, recruitment of C2 to the PIDDosome and the resulting induced proximity leads to enforced association of the two Venus halves, culminating in the C2 BiFC signal (see Figure 4K). Thus, Venus fluorescence acts as a terminal readout for PIDDosome assembly (Bouchier-Hayes et al., 2009). HeLa.C2 Pro-BiFC cells (Ando et al.,

2017) harbor a bicistronic construct in which C2 Pro-VC and C2 Pro-VN, separated by the viral 2A self-cleaving peptide, are translated from a single mRNA transcript. This ensures that C2 Pro-VC and C2 Pro-VN are expressed at equal levels. The sensitivity of the endogenous C2 BiFC reporter is sufficient to detect C2 induced proximity in the nucleolus, a major but not unique site for PIDDosome formation (Ando et al., 2017). Parental HeLa.C2 Pro-BiFC cells (0.5 x $10^5$ cells) (Ando et al., 2017) were seeded directly on coverslips, transfected with siRNAs for 48 hours and treated with qVD-OPH (20 μM) and IR (10 Gy) and harvested 24 hours post-treatment. For the experiments including overexpression of IRAK1 cDNA, DNA transfections were done 24 hours after siRNA transfection, treated with qVD-OPH (20 μM) and IR (10 Gy) and harvested 24 hours post-treatment. Cells expressing the BiFC components were identified by fluorescence of the linked mCherry protein in stable cell lines. Venus channel image data was analyzed to determine the cells positive for C2 BiFC. More than 100 cells were counted over three independent experiments.

**Two-photon laser micro-irradiation of SV40 MEF**

SV40-immortalized MEFs were transfected with 1.5μg of either pDsRed-PCNA or pRFP-Ku70 expression plasmids using Lipofectamine 2000 (Invitrogen) according to the manufacturer’s instructions. The next day, ~$10^4$ transfected cells were seeded into a 35mm glass-bottom dish and treated with 20μM BrdU. Live-cell imaging was performed 24 hours after BrdU treatment on a Nikon Ti Eclipse inverted microscope (Nikon Inc, Tokyo, Japan) equipped with A1 RMP (Nikon Inc) confocal microscope system (Nikon Inc) and Lu-N3 Laser Units (Nikon Inc). Laser micro-irradiation and imaging were conducted via the NIS Element High Content Analysis software (Nikon Inc) using a 800nm 2-photon laser (10μm x 0.5μm rectangular region, energy level ≅ 2800mW). Images of each cell of interest were acquired at 60x magnification immediately before and immediately after micro-irradiation (~2 seconds acquisition time), as well as 5 minutes after micro-irradiation. 15-20 minutes after micro-irradiation, cells were washed twice with ice-cold PBS, and then fixed in 1% paraformaldehyde, 0.25% Triton fixation buffer on ice for at least 30 minutes. The fixation buffer was replaced with PBS and cells were stored in 4°C until processed for IRAK1pT209 IF analysis.

**Zebrafish Lines, Maintenance and X-Ray Irradiation**

Adult zebrafish were maintained on a 14:10 hour light:dark cycle at 28°C in accordance with the regulations and policies of the Mount Sinai Institutional Animal Care and Use Committee. The study is compliant with all relevant ethical regulations regarding zebrafish research. The progeny of homozygous $p53^{M214K/M214K}$ ($p53^{MK/MK}$) fish were used throughout. M214 is orthologous to human M246, a commonly mutated residue within the mutational hotspot in the p53 DNA binding domain (http://www-p53.iarc.fr/) (Olivier et al., 2002; Olivier et al., 2010). $p53^{MK/MK}$ embryos demonstrate fully penetrant resistance to IR-induced cell death (Sidi et al., 2008), a phenotype suppressed by genetic or pharmacological targeting of IRAK1 (Liu et al., 2019). The TILLING-mediated generation of the $p53^{M214K/M214K}$ line, including allele designation, has been described (Berghmans et al., 2005; Sidi et al., 2008). X-ray irradiation of embryo clutches at 18 hours post-fertilization (hpf) was performed using the X-RAD 320 (PXI Precision X-ray, filter 2) at Mount Sinai hospital irradiator CoRE facility.

**Micro-injections into zebrafish embryos**

*Synthetic mRNAs* – Wild-type (and mutant, when indicated) full-length cDNAs for human IRAK1 (hIRAK1), human IRAK4 (hIRAK4), human Peli1 (hPeli1) and human Peli2 (hPeli2) were subcloned from parent mammalian expression vectors (see Plasmids and DNA transfection into cultured human cancer cells) into the zebrafish expression vector pCS2+. All plasmids were linearized by Sac2 single enzyme treatment at 37 $^{o}$C for 4 hrs. Digests were stopped by adding 1/20 volume 0.5 M EDTA, 1/10 volume of 3 M Na acetate and 2 volumes EtOH. Samples were mixed and chilled at -20 $^{o}$C for 15 min, washed and resuspended in TE buffer. Sense-capped mRNAs were synthesized for injection using the mMESSAGE mMACHINE SP6 kit (Ambion, #AM1340) following the manufacturer's instructions. mRNA concentrations were determined by Nanodrop and RNA gel. Approximately 20 pg of the synthetic human mRNAs were co-injected with MOs to *irak4*, *irak1*, *peli1b* or *peli2*, depending on the experiment, during the 1-2 cell stage. Importantly, these synthetic mRNAs encoding human proteins lacked the MO target sequences on the corresponding zebrafish endogenous mRNAs. This ruled out that any phenotypic rescue resulted from MO titration by the co-injected synthetic mRNA.

*Morpholino antisense oligonucleotides (MOs)* – Splice-junction MOs (Gene Tools LLC) to *irak1*, *irak4* and *myd88* have been previously described (Liu et al., 2019), and MOs targeted to *peli1b* and *peli2* were designed by and obtained from Gene Tools LLC. MOs were resuspended in sterile water to a stock concentration of 1 mM. Approximately 1 nl 0.25-1 mM MO was delivered into one-cell stage zebrafish embryos by microinjection. Synthetic mRNAs were in vitro transcribed from pCS2+ templates using the Ambion mMESSAGE mMACHINE kit according to the manufacturer's instructions, dispensed as 2 ml aliquots, and stored at -80 $^{o}$C. Synthetic mRNAs were diluted in RNase free water to a final concentration if 25 ng/ml. Approximately 1 nl mRNA was delivered into one-cell stage embryos by microinjection. All micro-injections of MOs or mRNAs were performed with a NARISHIGE IM 300 microinjector as described in detail online (bio-protocol.org/prep1293). MO-mediated gene knockdowns efficiencies were verified in whole-embryo extracts by western blot when possible or by RT-PCR to detect exon-skipping or intron-retention events (i.e., Irak4, Peli2). All MO sequences and RT-PCR primers are listed in Supplementary Table 1.

**RT-PCR and Protein Extraction from Zebrafish Embryos**

Embryonic RNA was isolated from 24-48 hpf embryos (>15 embryos/sample) using a standard Trizol method (250 µL Trizol (Invitrogen), 50 µL $CHCl_3$, 175 µL isopropanol). One microgram of purified RNA was used to generate cDNA using the Invitrogen SuperScript First Strand III RT-PCR kit, with oligo-dT primers. Two micrograms of the cDNA product were loaded on a 1% agarose gel. Pooled embryo protein lysates were harvested as previously described (Sidi et al., 2008) and analyzed by western blot (see Western blotting and Antibodies).

**Acridine Orange (AO) Labeling of live embryos.**

At 24 h post-IR, live embryos were labelled with acridine orange at 10 mg $ml^{-1}$ in egg water for 20 min, then dechorionated in pronase (2.0 mg $ml^{-1}$ in egg water) for 5 min and rinsed at least three times for 20 min in egg water. Embryos were imaged with a SMZ1500 fluorescent stereomicroscope and NIS-Element software (Nikon), and analyzed with ImageJ as previously described (Sidi et al., 2008).

**Whole-mount immunofluorescence imaging**

Zebrafish embryos were injected with 25 pg human IRAK1 mRNA at the one cell stage, irradiated at 18 hpf (see Zebrafish Lines and Maintenance) and fixed at 15 min post-IR in 4% PFA overnight at 4 $^{o}$C. Embryos were rinsed three times in PBST, dehydrated with 1:2, 2:1 methanol:PBST, then 3 times in methanol, 5 min each, and stored at -20 $^{o}$C overnight. Embryos were rehydrated with 2:1, 1:2 methanol:PBST, then 3 times PBST, 5 min each. Embryos were permeabilized with proteinase K (10 mg/ml in PBST) for 3 min, quickly washed 3 times with PBST and fixed with 4% PFA for 20 min. After 5 washes in PBST, 5 min each, 400 ml of blocking reagent was added (1% BSA + 10% normal goat serum (NGS, 10000C, invitrogen) in PBST) and embryos were stored at 4 $^{o}$C overnight. Anti-IRAK1pT209 was added at 1:100 final concentration in blocking reagent and embryos were incubated at 4 °C over 3 nights on a shaker. Embryos were washed 3 times with PBST, blocked again at 4 °C overnight, and secondary antibody was added (1:300) 4 °C overnight. Finally, embryos were washed 5 times with PBST and processed for whole-mount immunohistochemistry as follows as depicted in diagram below. Images were obtained under a 40x oil objective with an inverted confocal microscope (405 nm, 488 nm; SP5, Leica) and acquired using LAS software. The setting of the software was: size 1024X1024, zoomX2, pinhole 1AU, average 4 for AlexaFluor 555 channel and 2 for all other channels, with Z stacks of 1 mm/slice.

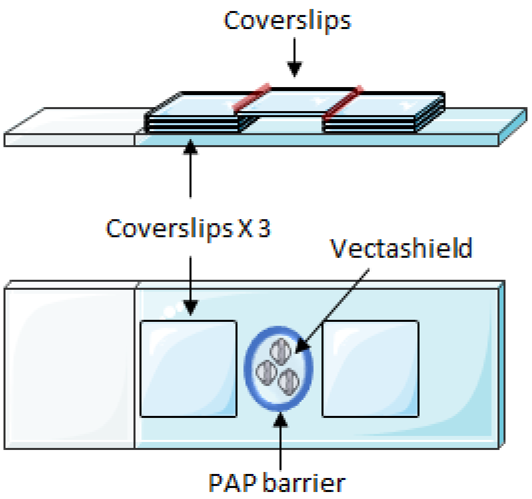

**Phylogenetic analyses**

Multiple sequence alignments of IRAK1 (Figure 4A) and Peli homologs (Figure S6E) across various species were performed using Clustal Omega (https://www.ebi.ac.uk/Tools/msa/clustalo/).

**Statistical Analysis**

Paired two-tailed Students t-tests were used to determine p-values ($\alpha = 0.05$). The log rank test was used to determine p-values for survival curves. Data in bar graphs are represented as mean ± SD or ± SEM, as indicated in legends, and statistical significance was expressed as follows: *, $P < 0.05$; **, $P < 0.01$; ***, $P < 0.001$; ns, not significant.

#### Supplementary Figure Legends

**Figure S1. Extended data for Main Figure 1.**

(A) HeLa, CAL27, MCF7 and MDA-MB-231 cells transfected with the indicated siRNAs were treated with or without IR (7.5 Gy) and stained with the vital dye alamarBlue 72 hr post-IR. Data are means +/− SD of 3 independent experiments, $*p < 0.05$, $**p < 0.01$, $***p < 0.001$, two-tailed Student's t-test.

(B) HeLa, CAL27, MCF7 and MDA-MB-231 cells transfected with the indicated siRNAs as in (A) were harvested 48 hours post-transfection and analyzed by western blot with indicated antibodies to assess knockdown efficiencies.

(C) Reconstituted *IRAK4* null cells from Main Figure 1C were analyzed by western blot to verify uniform expression of the indicated FLAG-IRAK4 variants.

(D) Individual channels for images shown in Main Figure 1I.

**Figure S2. Extended data for Main Figure 2.**

(A-B) HeLa cells treated with IR (7.5 Gy) were fixed at indicated time points (min post-IR), stained with anti-IRAK1 (A) or anti-IRAK1pT209 (B) antibodies in parallel, co-stained with DAPI and imaged by confocal microscopy. Note that while IRAK1pT209 gradually accumulates in the nucleus, native IRAK1 remains largely cytoplasmic.

(C-D) Indicated cell lines were treated with or without IR (7.5 Gy), fixed at 15 min-post IR, stained with anti-IRAK1pT209 antibody and DAPI, and imaged by confocal microscopy. Images representative of at least two independent experiments were quantified in (D).

(E) (Left) Schematic of lateral and dorsal views of the 18 hour zebrafish embryo, with imaged area boxed in pink. A, anterior; P, posterior; V, ventral; D, dorsal; L, left; R, right. (Right) Zebrafish *p53*$^{MK/MK}$ mutant embryos were co-injected at the 1 cell stage with *irak1* MO and synthetic mRNA for human FLAG-IRAK1 to detect IRAK1p209 in vivo (the IRAK1pT209 antibody does not cross-react with zebrafish Irak1). Embryos were treated with or without whole-body IR (12.5 Gy) 18 hours later, fixed at 15 min post-IR, and stained as whole-mounts with IRAK1pT209 antibody and DAPI. Higher magnification images of boxed areas shown to the right. Anterior and spinal cord areas delineated by thick and thin dashed lines, respectively. Scale bar, 40 μm.

(F) HeLa cells treated with IR (7.5 Gy) were fixed at indicated time points (hours post-IR), stained with anti-IRAK1pT209 and anti-γH2A.X antibodies, co-stained with WGA and DAPI and imaged by confocal microscopy. Images from 3 independent experiments were quantified in (G).

(G) Quantification of IRAK1pT209 stains such as in (F). Data are means +/− SD of 3 independent experiments. Statistical significance vs. bar 1: *$p$ < 0.05, **$p$ < 0.01, ***$p$ < 0.001; two-tailed Student's t-test.

(H) Parental and *IRAK4*$^{-/-}$ HeLa cells were transfected with or without the indicated FLAG-IRAK4 constructs (WT, D329A or R12C), treated with or without IR (7.5 Gy), fixed at 15 min post-IR, stained with anti-IRAK1pT209 antibody and DAPI, and imaged by confocal microscopy. Images representative of two independent experiments. WT, wild-type; D329A, kinase-dead; R12C, variant unable to bind MyD88.

(I) Cells as in (H) were lysed at 15 min post-IR and analyzed by western blot with indicated antibodies. Note that the western confirms the IF results in (H), whereby WT and R12C, but not D329A, FLAG-IRAK4 restore IRAK1 phosphorylation on T209 in irradiated *IRAK4*$^{-/-}$ cells.

**Figure S3. Extended data for Main Figure 3.**

(A) HeLa, CAL27, T98G, MDA-MB-231, MCF7 and SAS cells were treated with or without IR (7.5 Gy), fixed at 15 min post-IR, stained with anti-IRAK1pT387 antibody and DAPI, and imaged by confocal microscopy. Images representative of two independent experiments. HNSCC, head and neck cancer; GBM, glioblastoma multiforme; BrCA, breast cancer.

(B) Cells from Main Figure 3 (A-H) were lysed and analyzed by western blot with indicated antibodies to assess the expression levels of the transfected FLAG-IRAK1 constructs.

(C-D') HeLa cells grown on cover slips were treated with or without IR (7.5 Gy), fixed at 5 or 15 min post-IR, stained with IRAK1pT387 (C,D) or IRAK1pT209 (C',D') and nucleolin (NCL, (C,C')) or fibrillarin (D,D') antibodies, co-stained with DAPI and imaged by confocal microscopy. Note that IRAK1pT387, but not IRAK1pT209, colocalizes with nucleolar markers at 15 min post-IR. Higher magnification views of boxed areas in C and C' shown in Main Figure 3K.

**Figure S4. Extended data for Main Figure 4.**

(A) HeLa cells transfected with the indicated siRNAs were treated with or without IR (7.5 Gy), fixed 15 min post-IR, stained with IRAK1pT209 antibody and DAPI, and imaged by confocal microscopy. Relative cytoplasmic (left) and nuclear (right) IRAK1pT209 signals were quantified from 3 independent experiments. Data are means +/− SD, with statistical significance vs. bar 2: **$p<0.01$, ***$p < 0.001$; two-tailed Student's t-test.

(B) Representative images from the experiments analyzed in (A).

(C) Full western blot for Main Figure 4D to show non-irradiated cells.

(D) An additional experiment as shown in Main Figure 4B. Parental and non-irradiated controls are also included for comparison with reconstituted *IRAK1*$^{-/-}$ cells. Data were quantified in Main Figure 4C.

**Figure S5. Extended data for Main Figure 5.**

(A) HeLa cells transfected with the indicated siRNAs were treated with or without IR (7.5 Gy), fixed 15 min post-IR, stained with IRAK1pT209 antibody and DAPI, and imaged by confocal microscopy. Images from 3 independent experiments were quantified in (B-C). Note that siTLR7 and siTLR8 were co-transfected to address functional redundancy.

(B-C) Images from experiments as in (A) were analyzed for both percentage of IRAK1pT209 positive cells (B) and total signal intensity (C). Data are means +/− SD from 3 independent experiments, with statistical significance vs. bar 2: ns, non-significant, ***$p < 0.001$; two-tailed Student's t-test.

(D) Whole cell lysates from an experiment as in (A) were analyzed by western blot with indicated antibodies to verify knockdown efficiency of the indicated siRNAs.

(E) HeLa cells treated with indicated doses of bleomycin were fixed at 15 min, stained with IRAK1pT209 antibody and DAPI and imaged by confocal microscopy. Images representative of 2 independent experiments.

(F-H) HeLa cells treated with camptothecin (1 μM) (F), topotecan (1 μM) (G) or aphidicolin (0.2 mM) (H) were fixed at indicated time points and stained with indicated antibodies and DAPI. Note that IRAK1pT209 and γH2A.X signals both strictly correlate (in time and intensity) and spatially overlap. Images representative of 2 independent experiments. 3 independent sets of images for the 6 hour time point (such as also shown in Main Figure 5E) were quantified in Main Figure 5F-J.

(I) RFP-Ku80 SV40 MEFs were treated with 2-photon laser micro-irradiation and stained with IRAK1pT209 antibody. Cells showing one or more Ku80+ foci were analyzed. Images representative of 17 analyzed cells over 2 independent experiments are shown. From these, a total of 27 RFP-Ku80+ foci were counted, of which 22 (81.5%) were positive for IRAK1pT209.

**Figure S6. Extended data for Main Figure 6.**

(A) HeLa cells transfected with the indicated siRNAs, including 3 independent siRNAs targeted to UNC5CL, were treated with or without IR (7.5 Gy), fixed 15 min post-IR, stained with

IRAK1pT209 antibody and DAPI, and analyzed by confocal microscopy. Images representative of 2 independent experiments.

(B) Cells from an experiment such as in (A) were lysed and analyzed by western blot with indicated antibodies. Each siRNA was effective at depleting UNC5CL.

(C) HeLa, MDA-MB-231 and SAS cells transfected with the indicated siRNAs were treated with or without IR (7.5 Gy), fixed 15 min post-IR, stained with IRAK1pT209 antibody and DAPI, and analyzed by confocal microscopy. Images representative of 2 independent experiments.

(D) Representative confocal images for the 3 independent experiments analyzed in Main Figure 6I.

(E) Clustal W multiple alignment-based phylogenetic tree of *C elegans* (ce), *Drosophila melanogaster* (dm), zebrafish (z), xenopus (x), mouse (m) and human (h) Pellino family members. zPeli1b was identified as most closely related to other vertebrate Peli1 proteins and was thus selected for MO-targeting studies.

(F) Pooled whole-embryo lysates from embryos injected with the indicated MOs were analyzed by western blot with indicated antibodies, demonstrating knockdown efficacy.

(G) Pooled whole-embryo RNA extracts from embryos injected with the indicated MOs were analyzed by RT-PCR for *peli2* and loading control *rppo*. Sequencing showed that the *peli2* MO, which targets the exon 3/intron 3 splice donor site, led to skipping of exon 3 (∆exon 3), resulting in an early truncation of endogenous Peli2.

**Figure S7. Extended data for Main Figure 7.**

(A-B) HeLa cells transfected with indicated GFP-Peli expression constructs were treated with IR (7.5 Gy), harvested at 15 min post-IR, immunoprecipitated with IRAK1 (A) or GFP (B) antibodies and analyzed by western blot.

(C) HeLa cells depleted of (left) or inactivated for (right) Peli1 and Peli2 were analyzed by western blot with the "anti-Pellino1/2" antibody (clone F7, see Methods). The Pellino1/2 antibody recognizes Peli1 but not Peli2 and was used to immunoprecipitate endogenous Peli1 in main Figure 7F-H.

(D) *PELI1*$^{-/-}$ cells transfected with indicated GFP-Peli1 expression constructs (see Main Figure 7A) were irradiated (7.5 Gy), harvested at 15 min post-IR, and immunoprecipitated with IRAK1 antibody. Pulldowns and whole-cell lysates were analyzed by western blot with indicated antibodies. All Peli1 variants retain the ability to interact with IRAK1 with the notable exception of Peli1$^{C}$.

(E) Model for the canonical and non-canonical IRAK signaling pathways (see main text).

#### Supplemental Table 1. Antibodies and sgRNA, siRNA and MO sequences

##### Antibodies for immunofluorescence

| | | FIXATION | PERMEABILIZATION | Primary Antibody | Secondary Antibody |
|---|---|---|---|---|---|
| Active Caspase3 | BD Pharmingen | 1% PFA | 0.25% Triton(in 1X PBS) | 1:100 in 1% BSA(in 1X PBS) | 1:200 in 1% BSA (in 1X PBS) |
| Fibrillarin mAb | SCBT | | 0.25% Triton(in 1X PBS) | 1:200 in 1% BSA(in 1X PBS) | 1:200 in 1% BSA (in 1X PBS) |
| IRAK-1 (C-2) mAb | SCBT | 1% PFA | 0.25% Triton(in 1X PBS) | 1:100 in 1% BSA(in 1X PBS) | 1:200 in 1% BSA (in 1X PBS) |
| IRAK1 (D51G7) Rabbit mAb | CST | 1% PFA | 0.25% Triton(in 1X PBS) | 1:100 in 1% BSA(in 1X PBS) | 1:200 in 1% BSA (in 1X PBS) |
| IRAK1 (Phospho-Thr209) pAb | Assay Biotechnology | 1% PFA | 0.25% Triton(in 1X PBS) | 1:100 in 1% BSA(in 1X PBS) | 1:200 in 1% BSA (in 1X PBS) |
| IRAK1 (Phospho-Thr387) pAb | Assay Biotechnology | 1% PFA | 0.25% Triton(in 1X PBS) | 1:100 in 1% BSA(in 1X PBS) | 1:200 in 1% BSA (in 1X PBS) |
| IRAK4 | CST | 1% PFA | 0.25% Triton(in 1X PBS) | 1:100 in 1% BSA(in 1X PBS) | 1:200 in 1% BSA (in 1X PBS) |
| Nucleolin C23 (D-6) mAb | SCBT | 1% PFA | 0.25% Triton(in 1X PBS) | 1:1000 in 1% BSA(in 1X PBS) | 1:200 in 1% BSA (in 1X PBS) |
| phospho-Histone H2A.X (Ser139) Antibody, clone JBW301 | Millipore Sigma | 1% PFA | 0.25% Triton(in 1X PBS) | 1:10000 in 1% BSA(in 1X PBS) | 1:200 in 1% BSA (in 1X PBS) |
| Phospho-IRAK4 (Thr345/Ser346) (D6D7) Rabbit mAb | CST | 1% PFA | 0.25% Triton(in 1X PBS) | 1:100 in 1% BSA(in 1X PBS) | 1:200 in 1% BSA (in 1X PBS) |
| Wheat Germ Agglutinin (WGA) CF555 | Biotium | | | 2.5:1000 in 1X PBS | |

**Antibodies for western blotting**

| Antibodies | Provider | $1^0$ Dilutions | $2^0$ Dilutions |
|---|---|---|---|
| ATM (D2E2) mAb | CST | 1:1000 in 5% BSA(TBS-Tween) | 1:5000 in 5% Milk (TBS-tween) |
| ATR pAb | CST | 1:1000 in 5% BSA(TBS-Tween) | 1:5000 in 5% Milk (TBS-tween) |
| FLAG (DYKDDDDK) pAb | CST | 1:1000 in 5% BSA(TBS-Tween) | 1:5000 in 5% Milk (TBS-tween) |
| GAPDH pAb (14C10) | CST | 1:5000 in 5% BSA(TBS-Tween) | 1:5000 in 5% Milk (TBS-tween) |
| GFP mAb( Clone 7.1 and 13.1) | Roche | 1:2000 in 5% BSA(TBS-Tween) | 1:5000 in 5% Milk (TBS-tween) |
| IRAK1 (D51G7) Rabbit mAb | CST | 1:1000 in 5% BSA(TBS-Tween) | 1:5000 in 5% Milk (TBS-tween) |
| IRAK-1 (C-2) mAb | SCBT | 1:1000 in 5% BSA(TBS-Tween) | 1:5000 in 5% Milk (TBS-tween) |
| IRAK4 | CST | 1:1000 in 5% BSA(TBS-Tween) | 1:5000 in 5% Milk (TBS-tween) |
| MyD88 (D80F5) Rabbit mAb | CST | 1:1000 in 5% BSA(TBS-Tween) | 1:5000 in 5% Milk (TBS-tween) |
| Pellino 1/2 (F-7) mAb | SCBT | 1:1000 in 5% BSA(TBS-Tween) | 1:5000 in 5% Milk (TBS-tween) |
| Pellino2 pAb Cat# NBP2-88033 (N-term) | Novus | 1:1000 in 5% BSA(TBS-Tween) | 1:5000 in 5% Milk (TBS-tween) |
| Pellino2 pAb Cat# H00057161-B01P (FL) | Novus | 1:1000 in 5% BSA(TBS-Tween) | 1:5000 in 5% Milk (TBS-tween) |
| Pellino2 pAb (C-term) 281-420 aa | Proteintech | 1:1000 in 5% BSA(TBS-Tween) | 1:5000 in 5% Milk (TBS-tween) |
| IRAK1 (Phospho-Thr209) pAb | Assay Biotechnology | 1:1000 in 5% BSA(TBS-Tween) | 1:5000 in 5% BSA (TBS-tween) |
| IRAK1 (Phospho-Thr387) pAb | Assay Biotechnology | 1:1000 in 5% BSA(TBS-Tween) | 1:5000 in 5% BSA (TBS-tween) |
| Phospho-IRAK4 (Thr345/Ser346) (D6D7) Rabbit mAb | CST | 1:1000 in 5% BSA(TBS-Tween) | 1:5000 in 5% BSA (TBS-tween) |
| TLR3 (TLR3.7) mAB | SCBT | 1:1000 in 5% BSA(TBS-Tween) | 1:5000 in 5% Milk (TBS-tween) |
| TLR4 (25) mAB | SCBT | 1:1000 in 5% BSA(TBS-Tween) | 1:5000 in 5% Milk (TBS-tween) |
| TLR7 (4F4) mAb | SCBT | 1:1000 in 5% BSA(TBS-Tween) | 1:5000 in 5% Milk (TBS-tween) |
| TLR8 (D-8) mAb | SCBT | 1:1000 in 5% BSA(TBS-Tween) | 1:5000 in 5% Milk (TBS-tween) |
| UNC5CL (AT116) pAb | Enzo Life sciences | 1:500 in 5% BSA(TBS-Tween) | 1:5000 in 5% Milk (TBS-tween) |

**siRNA sequences**

| siRNA | Provider | Sequence |
|---|---|---|
| siATM | Qiagen | AAGGCTATTCAGTGTGCGAGA |
| siATR | Qiagen | CAGGCACTAATTGTTCTTCAA |
| siImportin B | Qiagen | AAGTACAATTGTTACAATAA |
| siImportin A | Qiagen | ACGAATTGGCATGGTGGTGAA |
| siIRAK1 | Qiagen | CCGGGCAATTCAGTTTCTACA |
| siIRAK4 | Qiagen | AACACCGTGAACCTCAGTTAT |
| siLACZ | Qiagen | AACGTAGCGGAATACTTCGA |
| siMYD88 | Qiagen | AACTGGAACAGACAAACTATC |
| siPellino1 | Qiagen | CAGGACTACATTATAAATTTA |
| siPellino2 | Qiagen | CACGTGATATAACTGGTTATA |
| siTLR3 | Qiagen | AAGAACTGGATATCTTTGCCA |
| siTLR4 | Qiagen | ACCATTGAAGAATTCCGATAA |
| siTLR7 | Qiagen | CAGCTGGGTATAAATTCATGA |
| siTLR8 | Qiagen | TAGGTGTTCACCAGAGACATA |
| siTLR9 | Qiagen | TGCCTTCGTGGTCTTCGACAA |
| siUNC5CL #6 | Qiagen | TTCGATGGCACTGCCCTAGAA |
| siUNC5CL #7 | Qiagen | TGCGGATGTTATTGGAGCCAA |
| siUNC5CL #8 | Qiagen | CAGGGCTACTCTAGGAATGGA |

**sgRNA sequences**

| | | |
|---|---|---|
| IRAK1 #1 | CACCGACACGGTGTATGCTGTGAAG | AAACCTTCACAGCATACACCGTGTC |
| IRAK1 #2 | CACCGAGGAGTACATCAAGACGGGA | AAACTCCCGTCTTGATGTACTCCTC |
| IRAK1 #3 | CACCGATTTATCCCACAGAAAGACC | AAACGGTCTTTCTGTGGGATAAATC |
| IRAK1 #4 | CACCGGATCAACCGCAACGCCCCGTG | AAACCACGGGCGTTGCGGTTGATCC |
| IRAK4 #1 | CACCGGATGAACGACCCATTTCTGT | AAACACAGAAATGGGTCGTTCATCC |
| IRAK4 #2 | CACCGGCCTCAATGTTGGACTAATT | AAACAATTAGTCCAACATTGAGGCC |
| IRAK4 #3 | CACCGGGTAGTGTATTAGCAGTTTT | AAACAAAACTGCTAATACACTACCC |
| IRAK4 #4 | CACCGGGCACCACAAATTGCACAGT | AAACACTGTGCAATTTGTGGTGCCC |
| Pellino1 | caccgGGCTCGTTAATTGACCTCTG | aaacCAGAGGTCAATTAACGAGCCc |
| Pellino2 #1 | caccgGGGCGTGGATATCACATGGA | aaacTCCATGTGATATCCACGCCCc |
| Pellino2 #2 | caccgTCCACGAGGGGGCTTCACCG | aaacCGGTGAAGCCCCCTCGTGGAc |
| Pellino3#1 | caccgATGGTGAACTCATCGTCCTG | aaacCAGGACGATGAGTTCACCATc |
| Pellino3#2 | caccgGTAACAGACACGTCCCCTGG | aaacCCAGGGGACGTGTCTGTTACc |

**Morpholino sequences**

| | |
|---|---|
| standard control (std) | 5′-CCTCTTACCTCAGTTACAATTTATA-3′ |
| irak1 | 5′-AATCCTGCAACACAACAGCCACATT-3′ |
| irak4 | 5′-GTGAACAGGTAAAGCCTCACAGGAT-3′ |
| myd88 | 5’-TCTTGACGGACTGGGAAACTCG -3’ |
| peli1b | 5’-TCCCGCTCCCAGTGTGAACGCAACA-3’ |
| peli2 | 5’-ACTGCCTGTCACAACACAACAAAAG-3’ |

### Li, Shah et al. Figure S1

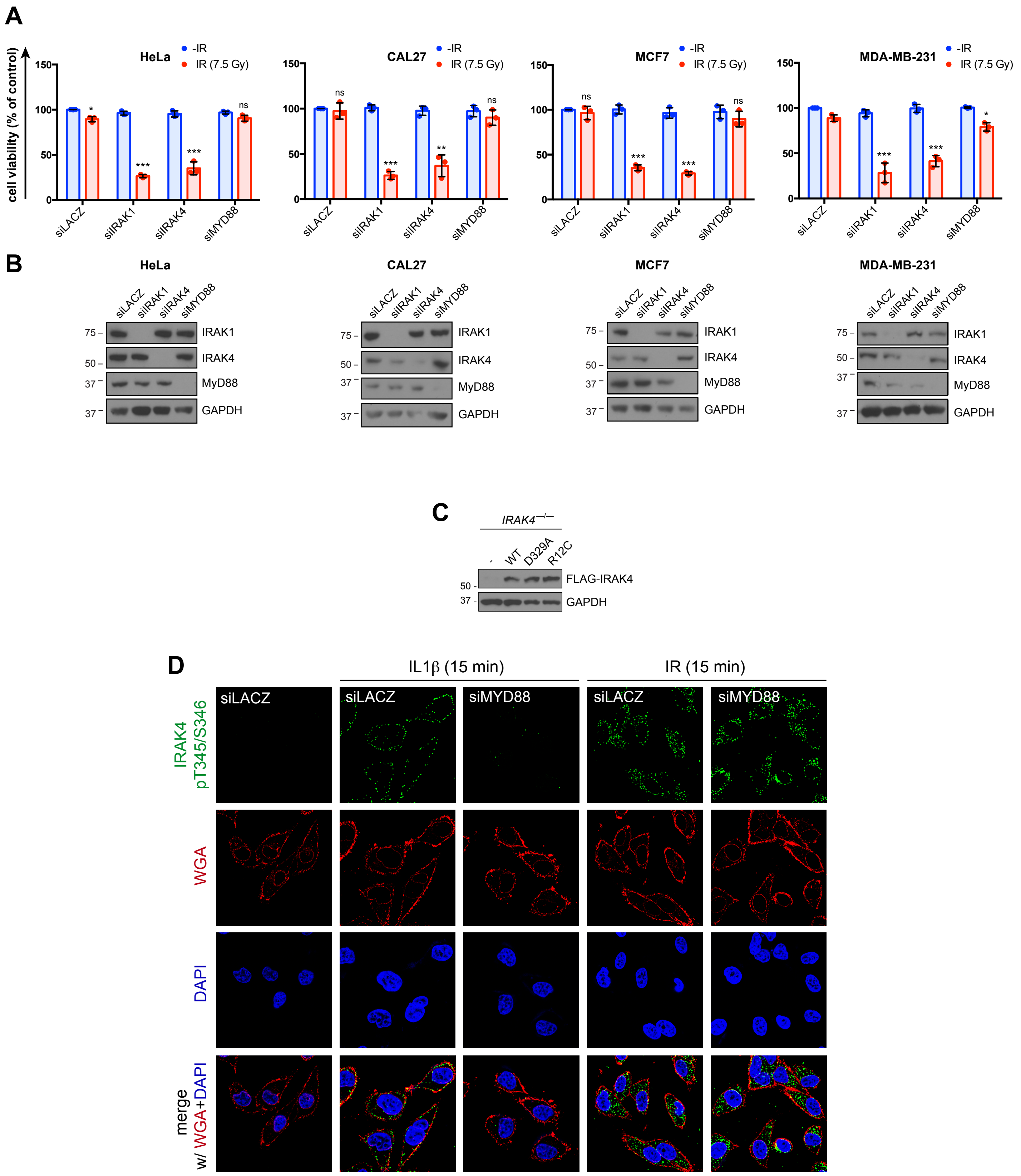

### Li, Shah et al. Figure S2

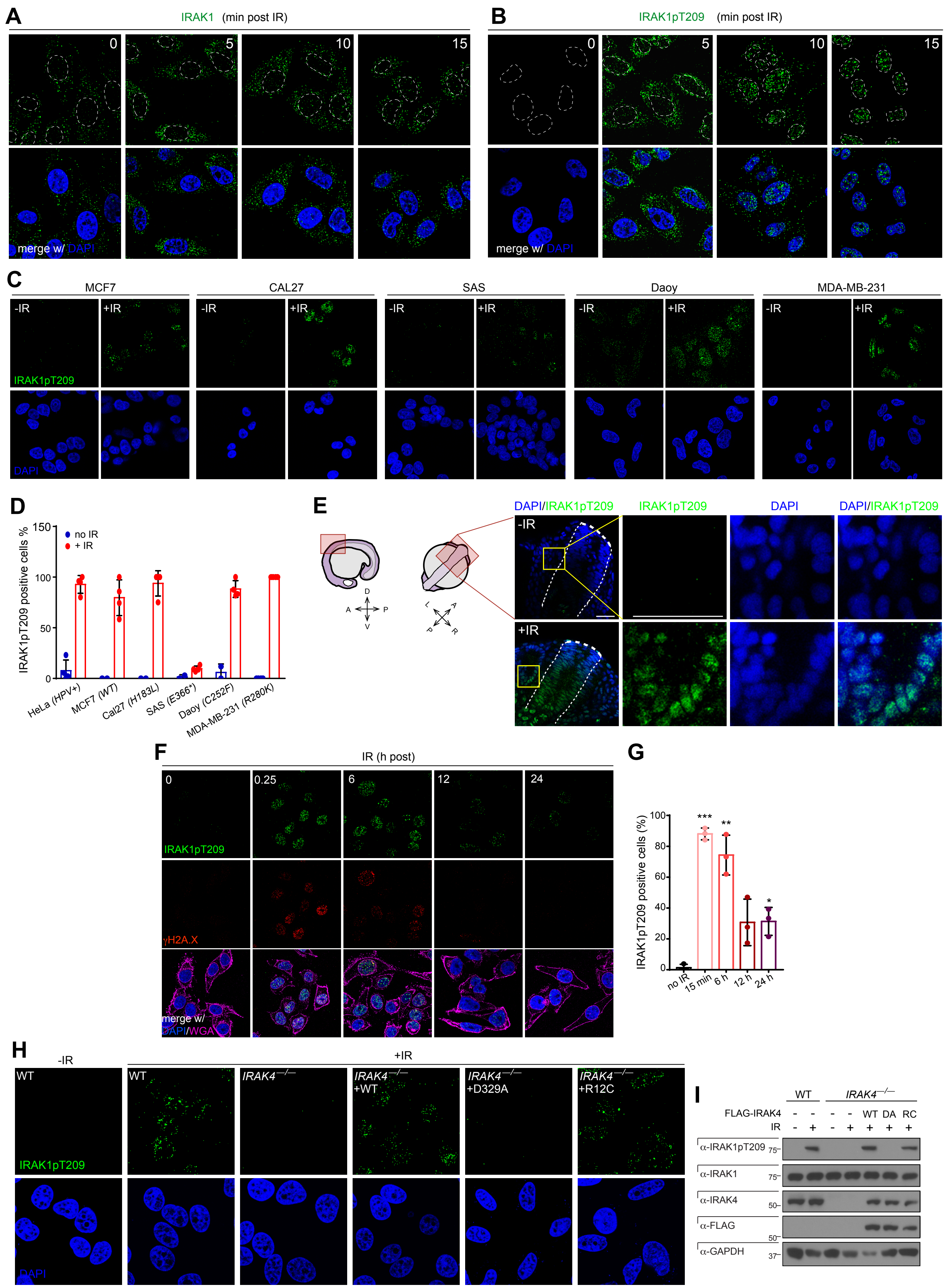

### Li, Shah et al. Figure S3

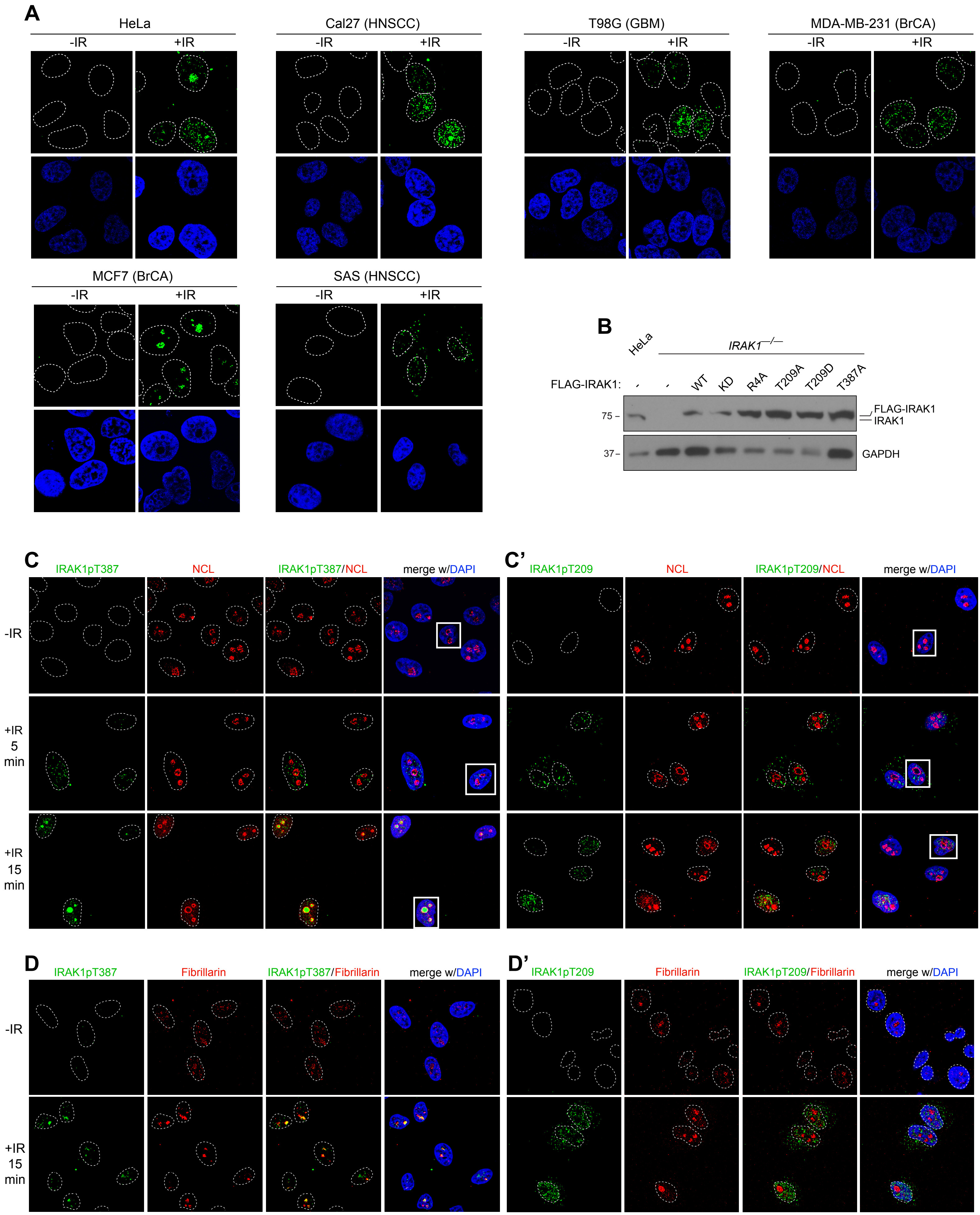

**Li, Shah et al. Figure S4**

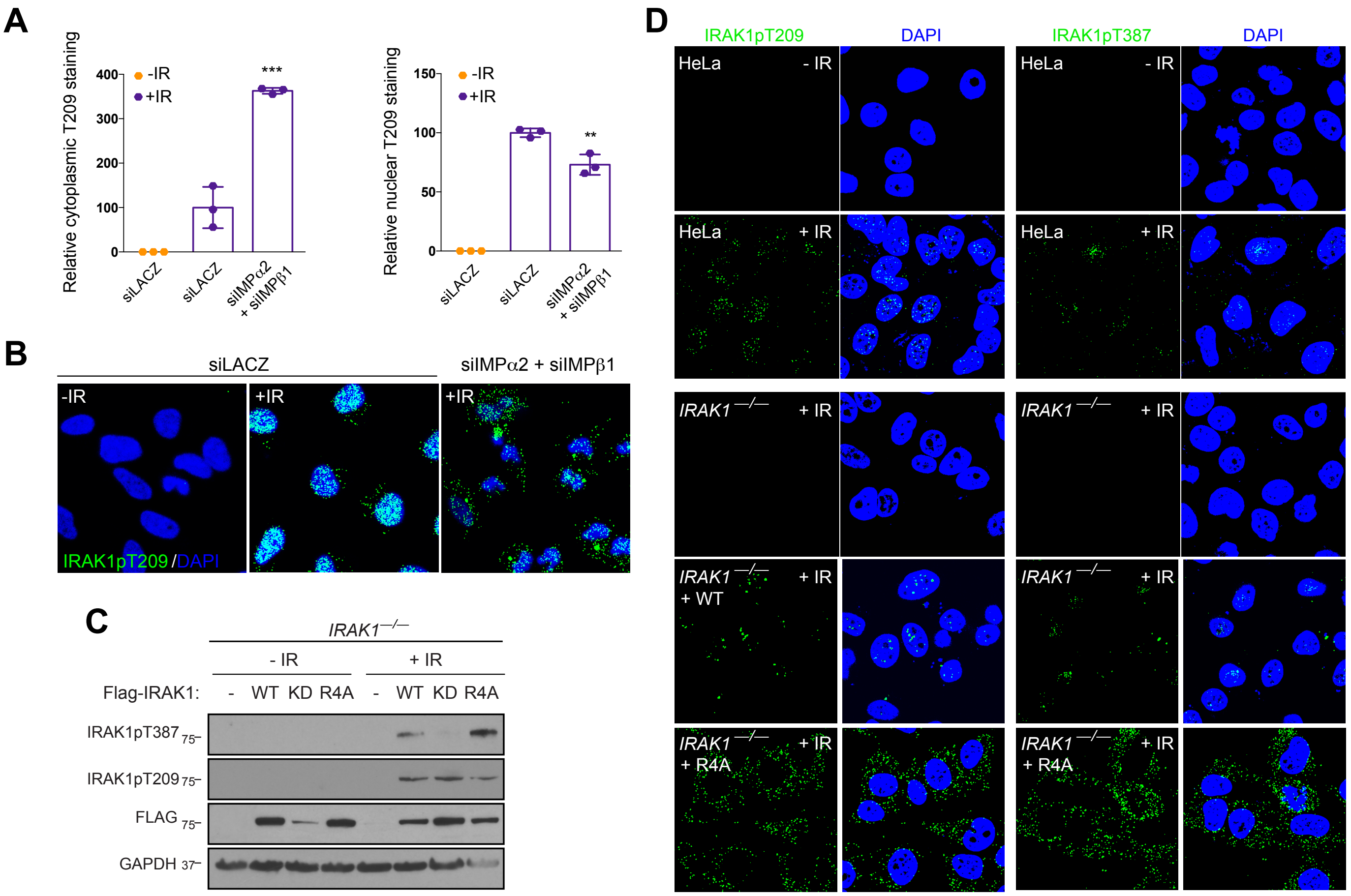

### Li, Shah et al. Figure S5

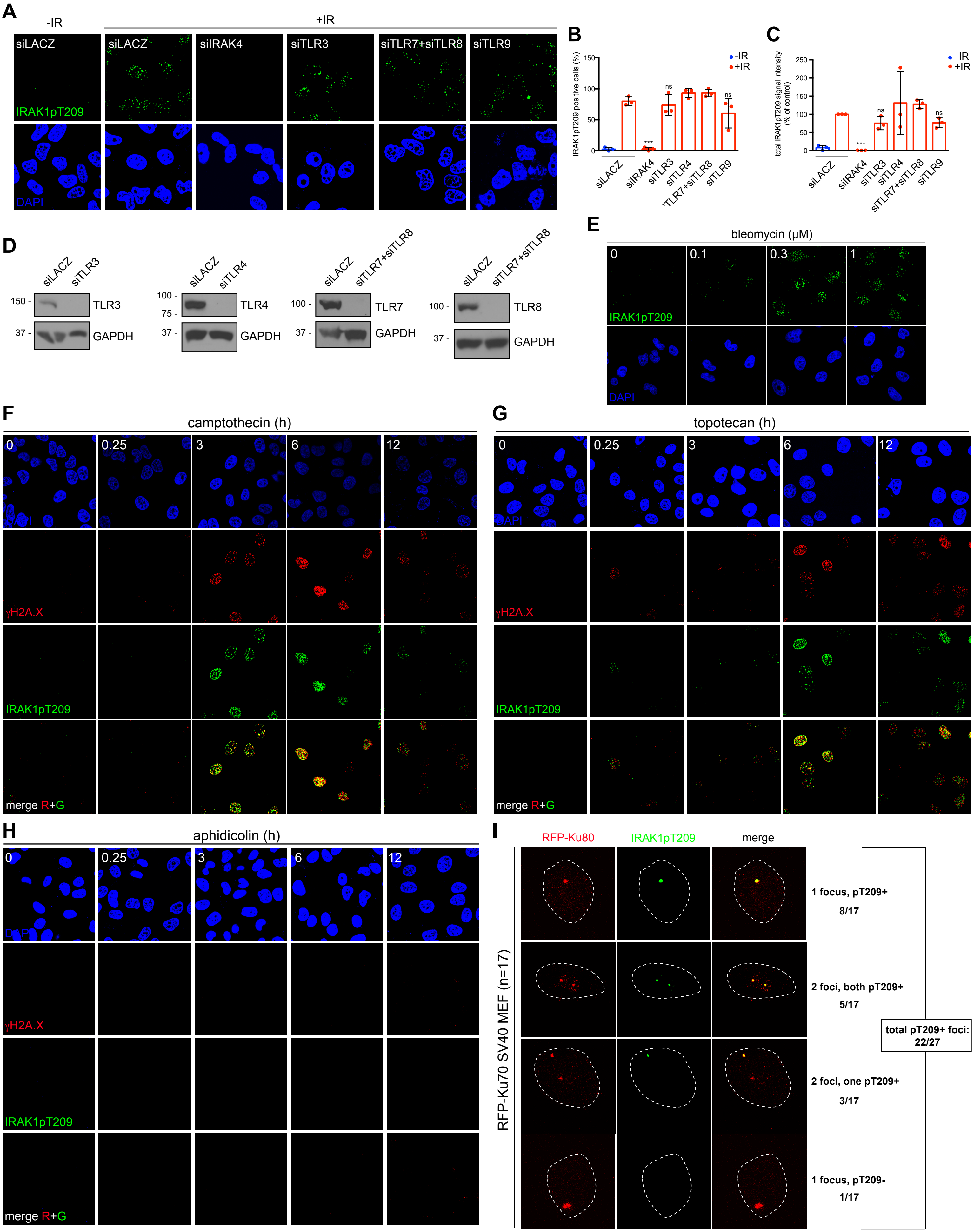

### Li, Shah et al. Figure S6

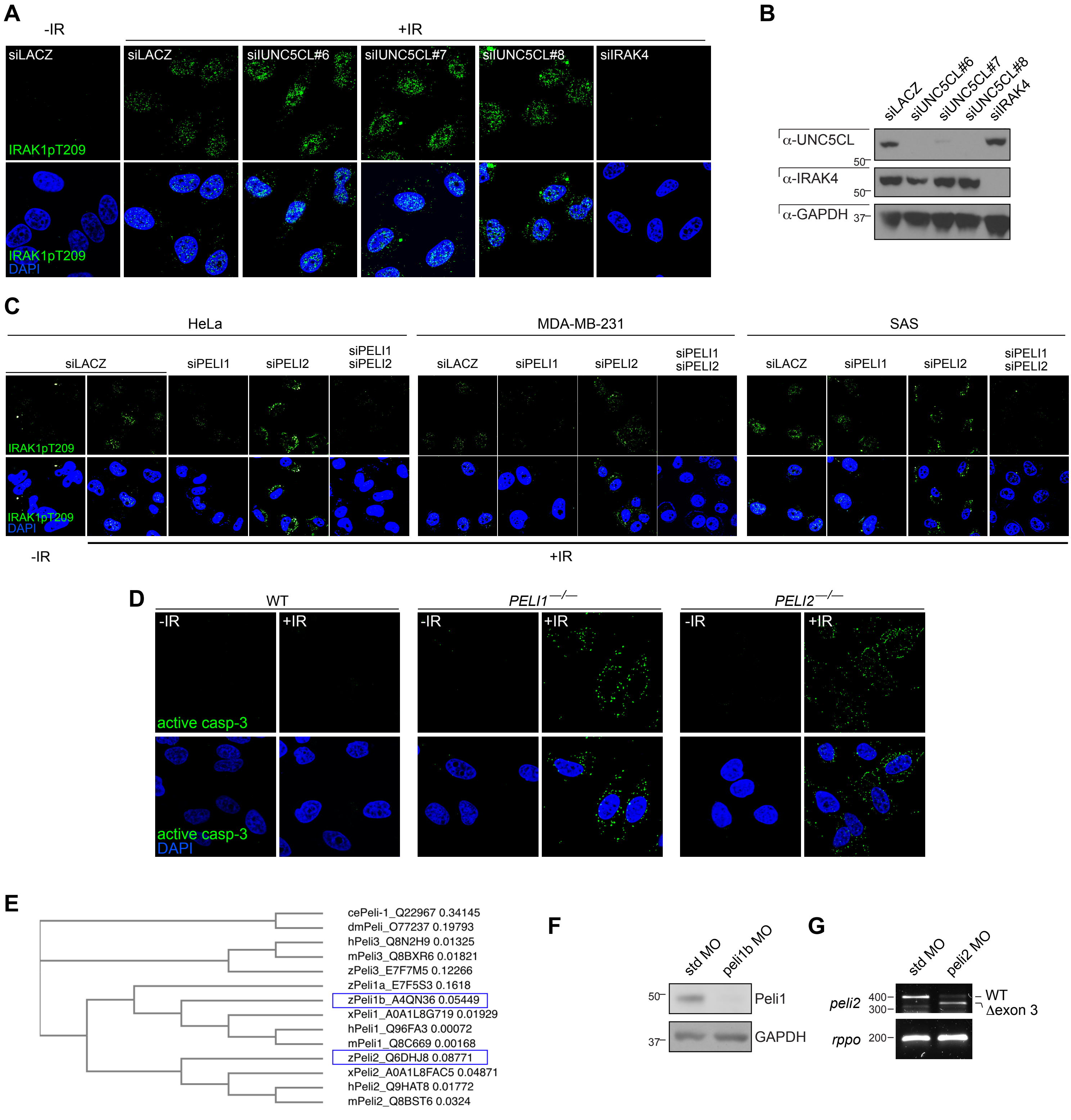

**Li, Shah et al. Figure S7**

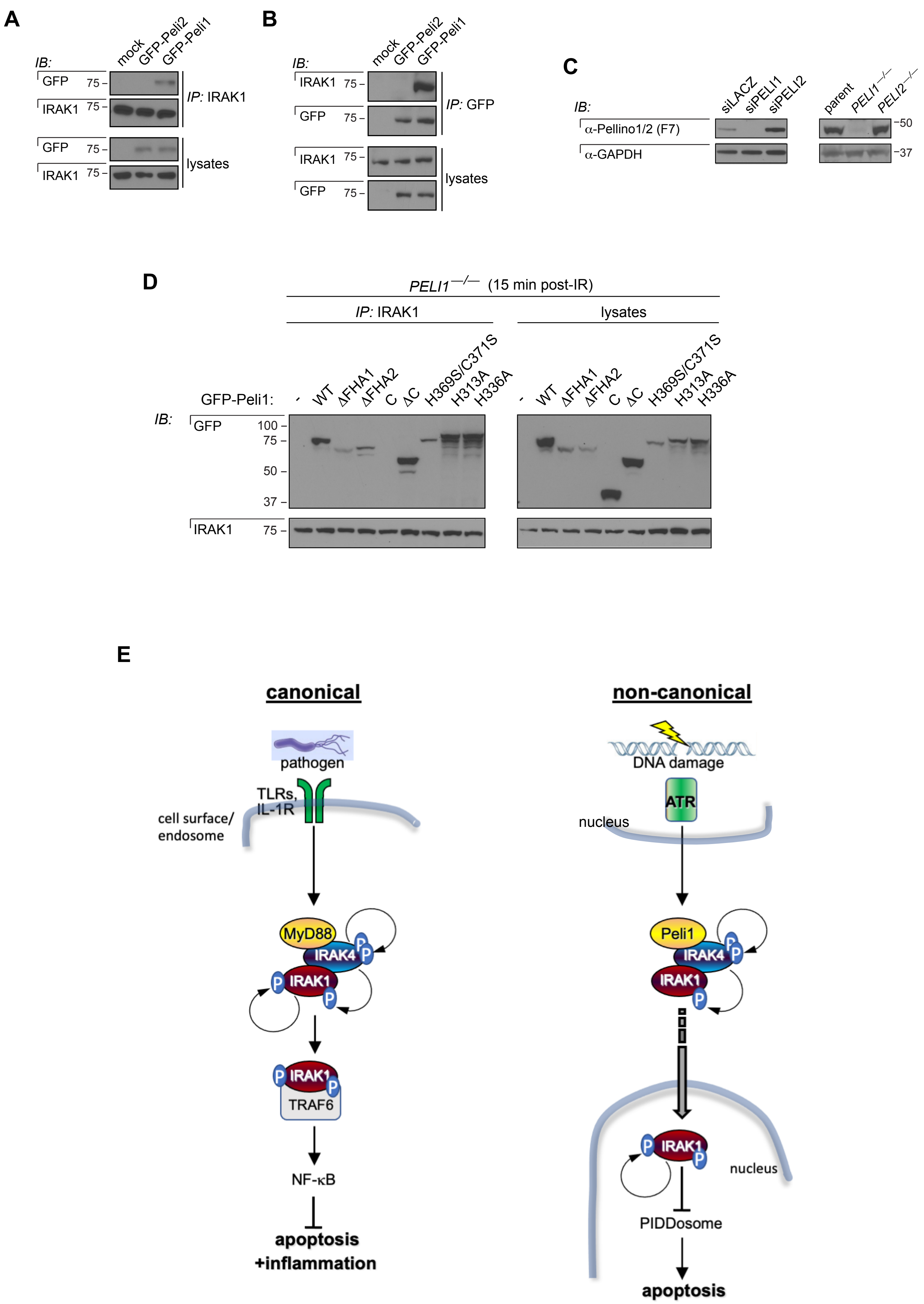
